# LET-381/FoxF and UNC-30/Pitx2 control the development of *C. elegans* mesodermal glia that regulate motor behavior

**DOI:** 10.1101/2023.10.23.563501

**Authors:** Nikolaos Stefanakis, Jessica Jiang, Yupu Liang, Shai Shaham

**Affiliations:** Laboratory of Developmental Genetics, The Rockefeller University, 1230 York Avenue, New York, NY 10065, USA; Research Bioinformatics, The Rockefeller University, 1230 York Avenue, New York, NY 10065, USA; Alexion Pharmceuticals, Boston, MA 02135, USA

**Keywords:** glia development, *let-381*, locomotory behavior, terminal selector, *unc-30*

## Abstract

While most CNS glia arise from neuroectodermal progenitors, some, like microglia, are mesodermally derived. To understand mesodermal glia development and function, we investigated *C. elegans* GLR glia, which ensheath the brain neuropil and separate it from the circulatory-system cavity. Transcriptome analysis suggests GLR glia merge astrocytic and endothelial characteristics relegated to separate cell types in vertebrates. Combined fate acquisition is orchestrated by LET-381/FoxF, a fate-specification/maintenance transcription factor expressed in glia and endothelia of other animals. Among LET-381/FoxF targets, UNC-30/Pitx2 transcription factor controls GLR glia morphology and represses alternative mesodermal fates. LET-381 and UNC-30 co-expression in naïve cells is sufficient for GLR glia gene expression. GLR glia inactivation by ablation or *let-381* mutation disrupts locomotory behavior and induces salt hypersensitivity, suggesting brain-neuropil activity dysregulation. Our studies uncover mechanisms of mesodermal glia development and show that like neurons, glia differentiation requires autoregulatory terminal selector genes that define and maintain the glial fate.

## INTRODUCTION

Glia are abundant and diverse cellular components of most, if not all, nervous systems, and are anatomically positioned to affect every aspect of signal transduction and processing in the brain. Glia dynamically regulate neuronal activity in response to presynaptic cues, provide insulation around axons and at synapses, and supply trophic support for neuron survival ^1^. Most glia, including astrocytes and myelinating glia, are derived from neuroectodermal precursors ^2,3^. By contrast, other glia, such as microglia, which are born in the yolk sac and migrate into the CNS, arise from mesodermal progenitors ^4–6^. Transcription factors regulating the development of some neuroectodermal glia are known ^2,7,8^; however, less is understood about the control of mesodermal glia differentiation. Furthermore, only a few factors sufficient to confer specification of certain glia subtypes in naïve cellular settings are known^9^.

The nematode *C. elegans* has been instrumental in uncovering basic principles of glia development and function ^10^. The nervous system of the adult *C. elegans* hermaphrodite contains 56 glial cells. 50 of these derive from the AB blast-cell lineage, which also produces neurons and epithelial cells. Six GLR glia derive from the MS blastomere, which primarily generates body-wall and pharyngeal muscle (**Fig. 1A**) ^11^. Thus, as in vertebrates, *C. elegans* possesses glia of both neuroectodermal and mesodermal origin. Some genes affecting *C. elegans* neuroepithelial glia development have been characterized ^12–15^; however, virtually nothing is known about GLR glia development or functions. GLR glia extend intricate, non-overlapping sheet-like processes that ensheath the inner aspect of the *C. elegans* brain neuropil (the nerve ring; **Fig. 1B**) and that are adjacent to neuromuscular synapses ^16^. At the nerve ring, GLR glia are electrically coupled to the RME motoneurons through gap junctions ^16^, and uptake extracellular GABA ^17^. More anteriorly, GLR glia extend thin processes that fasciculate with sensory neuron dendrites. GLR glia also physically separate the nerve ring from the pseudocoelomic body cavity which surrounds the pharynx and acts as a rudimentary circulatory system. The proximity to synapses and to the circulatory system, the association with GABA, and the ability to phagocytose injured neurons ^16,18,19^ make comparisons between GLR glia and astrocytes tempting.

**Figure 1.**
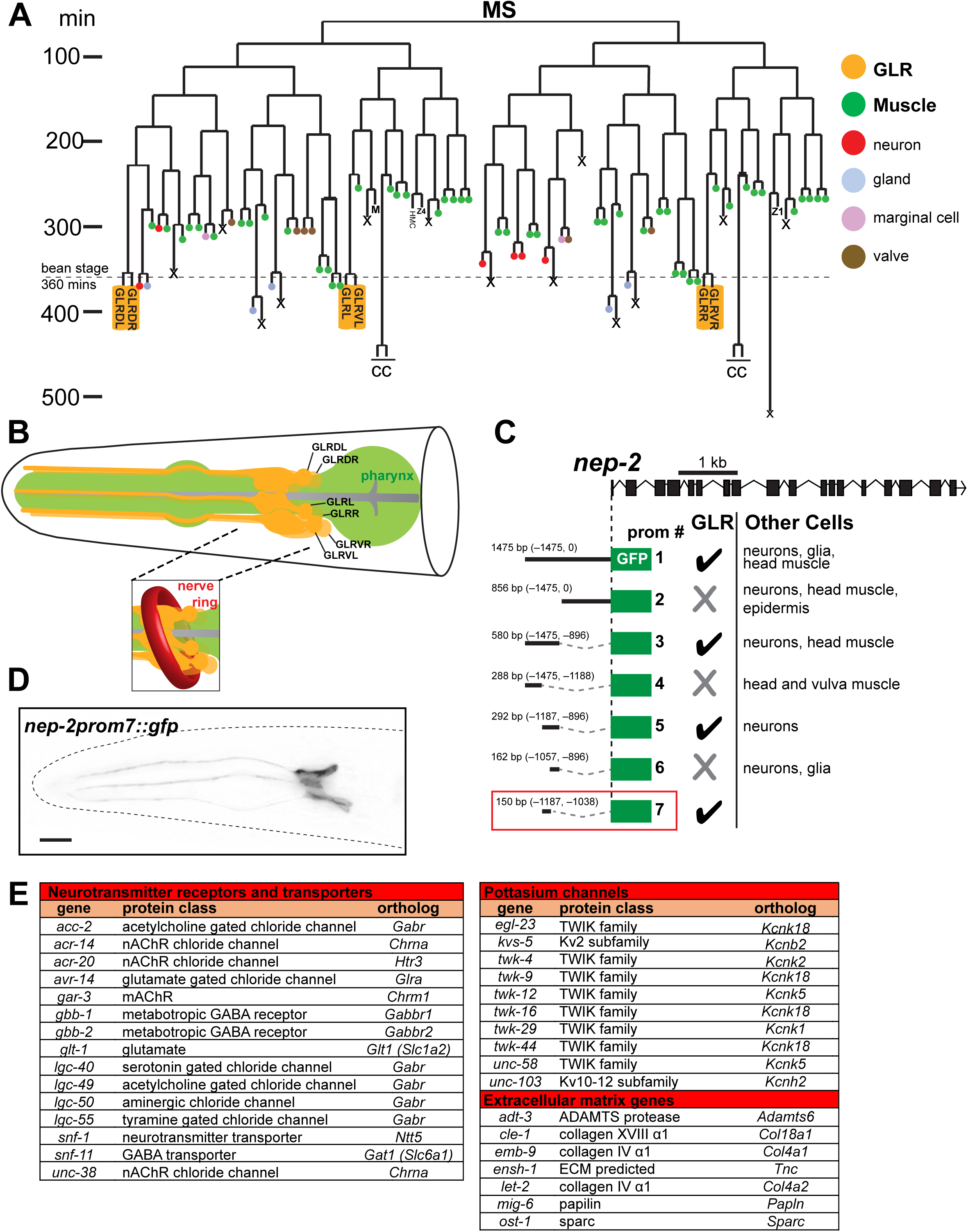
Generation of a GLR-specific driver to study the expression profile of the mesodermal GLR glia. (A) GLR glia (yellow boxes) derive from the lineage of the blast cell MS. This lineage produces mainly body wall muscle and pharyngeal muscle cells (green). GLR glia (yellow) are born at around the embryonic bean stage (360 minutes of embryonic development). The HMC cell and coelomocytes (CC) also derive from the MS lineage. Lineage schematic based on^11^. (B) Schematic representation of the GLR glia (yellow). Pharynx is shown in green. The inset shows how *C. elegans* Nerve Ring (red) wraps around the sheet-like GLR glia processes. Schematic drawn based on^19^. (C) Cis-regulatory dissection analysis for the gene *nep-2* resulted in isolation of a GLR glia-specific driver, prom7 (red box). (D) Fluorescence image of a L4 *C. elegans* showing expression of *nep-2prom7::gfp* specifically in GLR glia. Anterior is left, dorsal is up and scale bar is 10 μm. (E) Genes from three families (neurotransmitter receptors and transporters, potassium channels and extracellular matrix genes) are over-represented among GLR enriched genes.

Here, we describe the transcriptome of adult *C. elegans* GLR glia, uncovering similarities with both astrocytes and endothelial cells. We use the transcriptome to develop a molecular toolkit for labeling and manipulating GLR glia, which we use to identify and characterize two master regulators of GLR glia development. We show that LET-381, the *C. elegans* ortholog of the forkhead transcription factor FoxF, promotes GLR glia fate specification and maintenance; and that UNC-30/Pitx transcription factor controls GLR glia morphology and represses acquisition of an alternative mesodermal fate. Importantly, expression of both transcription factors in a naïve cell is sufficient to promote GLR glia gene expression. Through genetic studies, we order *let-381*, *unc-30*, and other genes into a pathway for GLR glia development. Finally, we show that *let-381* auto-regulation deficient mutants, as well as animals in which GLR glia are genetically ablated, exhibit specific defects in locomotory behavior and are hyper-sensitive to salt, suggesting important roles for these glia in coordinating neuronal activity.

## RESULTS

### GLR glia gene expression reveals similarities with astrocytes and endothelial cells

Although GLR glia reporters have been previously described, none are exclusively expressed in these cells ^20–24^. To identify drivers allowing specific genetic manipulation and marking of GLR glia, we performed promoter dissection studies of known GLR glia-expressed genes (**Appendix Figs. S1A-E**), isolating a 150 bp cis-regulatory sequence from the gene *nep-2* that, when fused to *gfp*, promotes expression only in GLR glia (**Figs. 1C and D**). Expression of this reporter is first detected in first-stage (L1) larvae and is maintained through adulthood.

To identify genes regulating GLR glia development and functions, we generated a stable transgenic *C. elegans* strain expressing nuclear YFP using the *nep-2* regulatory sequence (**Appendix Fig. S1F**). *nep-2prom7::nls::yfp* expressing cells were isolated from dissociated L4 larvae using fluorescence-activated cell sorting (FACS), and lysed to isolate mRNA. Following mRNA amplification and RNA-seq, we compared transcript abundances between GLR glia (YFP positive) and all other cells (YFP negative). We identified 886 genes with enriched GLR glia expression (p<0.05, log_2_-fold enrichment>1) out of 13794 genes with any GLR glia expression (>50 reads, **Appendix Fig. S1G**). To validate this list, we confirmed expression of 39 GLR glia-enriched genes using transgenic and endogenous *gfp* reporters. In addition, all previously known GLR glia genes show strong enrichment in our analysis.

Using PANTHER gene ontology over-representation analysis ^25,26^, we find that genes involved in synaptic transmission, including neurotransmitter receptors and transporters, potassium channels and genes encoding extracellular matrix proteins are over-represented among GLR glia-enriched genes (**Fig. 1E**). Genes from these transcript classes as well as other GLR enriched genes are over-represented in murine astrocytes (e.g. *snf-11/Gat1*, *gbb-1/Gabbr1*, *gbb-2/Gabbr2*, *glt-1/Glt1*, *ensh-1/Tnc, pll-1/Plcd4*) ^27–29^ suggesting similarities between GLR glia and astrocyte transcriptomes. Of note, ion, amino acid and neurotransmitter transporters, as well as extracellular matrix proteins are also enriched in endothelial cells of the blood brain barrier ^30^ while other GLR enriched genes, including *let-381/Foxf, dep-1/Ptprb, tag-68/Smad6, T16A9.4/Ece1, gei-1/Dlc-1, slcf-2/Slc2a1, unc-115/Ablim1*, *mrp-2/Abcc6*, are also enriched in endothelial cells of the central nervous system^28–30^. In vertebrates, astrocyte end-feet are found in proximity to endothelial blood vessels, and in *C. elegans*, GLR glia separate the circulatory cavity from the nerve ring. It is intriguing to speculate that to conserve cell numbers, *C. elegans* may have merged astrocytic and endothelial functions into the GLR glia cell type.

### *let-381/FoxF* is required early to specify GLR glia fate

To understand how the unusual fate merger of GLR glia arises, we sought to identify transcription factors that control GLR glia fate specification and differentiation. Transcripts encoding LET-381, the sole *C. elegans* ortholog of the Forkhead domain transcription factor FOXF ^31^, are highly enriched in GLR glia. Recent studies suggest that FoxF genes are expressed in phagocytic glia and independently in endothelial mural cells ^32,33^, raising the possibility that LET-381 could govern a combined gene expression pattern in GLR glia. To follow *let-381* expression in developing animals, we used CRISPR/Cas9 to insert *gfp* coding sequences into the endogenous *let-381* locus (**Fig. 2A**). Transgenic homozygotes display nuclear GFP fluorescence in GLR glia precursors (pre-bean; **Fig. 2B**) and in GLR glia until adulthood (**Fig. 2B**). Expression is also detected in the head mesodermal cell (HMC) and in coelomocytes, cell types also generated by the MS lineage (**Fig. 1A**). Animals carrying the *gfp* reporter transgene do not exhibit the lethality associated with loss of *let-381* (see below), suggesting that *let-381* gene function is retained.

**Figure 2.**
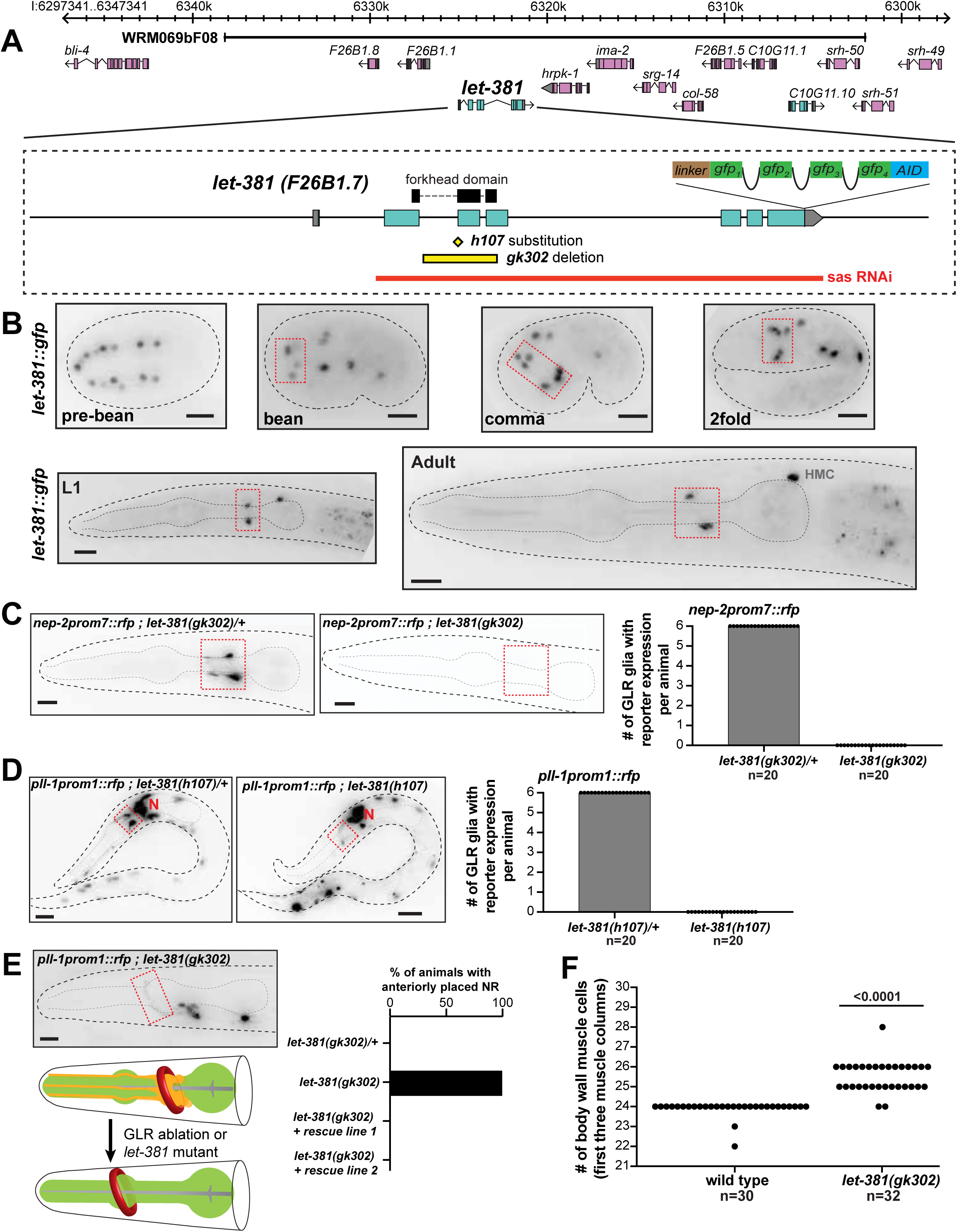
*let-381/FoxF* is required for GLR glia fate specification. (A) *let-381* genomic locus showing mutant alleles, reporters, fosmid genomic clones and RNAi sequences used in this study. (B) Expression of the endogenous *let-381::gfp* reporter at different stages during development. Dashed red boxes outline expression in GLR glia. (C) Absence of *nep-2prom7::rfp* expression in GLR glia (dashed red box) in homozygous *let-381(gk302)* mutants (right) as opposed to heterozygous animals (left). Quantification (number of GLR glia with *nep-2prom7::rfp* expression) is shown in the bar graph on the right. (D) Absence of *pll-1prom1::rfp* expression in GLR glia (dashed red box) in homozygous *let-381(h107)* mutants (right) as opposed to heterozygous animals (left). Quantification (number of GLR glia with *pll-1prom1::rfp* expression) is shown in the bar graph. Red “N” denotes *pll-1prom1::rfp* expression in neurons. (E) Similar to GLR glia-ablated animals (schematic), *let-381(gk302)* homozygous mutants lacking GLR glia exhibit anteriorly displaced nerve ring (NR). Dashed red box outlines a neuronal axon of the NR. Red circle in schematic indicates the Nerve Ring. Quantification is shown in the bar graph. Transgenic animals carrying the fosmid genomic clone WRM069bF08 with wild type *let-381* (rescue lines 1 and 2) display normal NR position. (F) Number of body wall muscle cells in the first three muscle columns of head and neck in wild type and *let-381(gk302)* mutants. Data information: Un-paired t-test was used for statistical analysis in (F). Anterior is left, dorsal is up and scale bars are 10 μm for all animal images.

To determine whether *let-381* promotes GLR glia fate specification, we introduced transgenic and endogenous GLR glia reporters into animals homozygous for the previously-identified *let-381* null alleles *gk302* or *h107*. *gk302* contains a deletion spanning LET-381 DNA binding domain encoding sequences; and *h107* is a G to A substitution at a splice acceptor site (**Fig. 2A**). Animals homozygous for either allele undergo late-embryonic/early-larval arrest, with a few *gk302* animals surviving to become sterile adults. Homozygous mutant *gk302* or *h107* animals fail to express five different GLR glia reporters that we tested (*nep-2prom7::gfp*, *pll-1prom1::rfp*, *gly-18prom::gfp*, *hlh- 1::gfp*, *unc-30::gfp*) (**Figs. 2C and D, Appendix Fig. S2A**), suggesting that GLR glia are not generated in these mutants. Consistent with this, laser ablation of GLR glia precursors was previously shown to cause anterior displacement of the nerve ring ^34^ and we find a similar defect in *let-381* mutants (**Fig. 2E**). In addition, GLR glia sister cells are body-wall muscle cells (**Fig. 1A**) and we observe extra muscle cells in the heads of *let-381(gk302)* mutants (**Fig. 2F**). All *let-381(gk302)* mutant defects are rescued by a transgene containing the wild-type *let-381* locus (WRM069bF08; **Figs. 2A and E, Appendix Figs. S2B and C**). Taken together, our results suggest that LET-381 is required for the generation of GLR glia, and in its absence, some presumptive GLR glia appear to acquire a default muscle cell fate.

### *let-381/FoxF* is continuously and cell-autonomously required to maintain GLR glia gene expression

Although LET-381 is required early to specify GLR glia, its expression during larval development and in adults suggests it may have additional later roles. To test this idea, we knocked down *let-381* in GLR glia of *let-381::gfp* animals by RNAi, using transgenic constructs co-transcribing sense and anti-sense *let-381* sequences ^35^ from the postembryonic *nep-2prom7* promoter. These animals, also homozygous for the RNAi-sensitizing allele *eri-1(mg366)*, down-regulate GFP expression specifically in GLR glia, but not in other *let-381* expressing cells, confirming RNAi specificity and efficacy (**Fig. 3A**). Importantly, *let-381* RNAi transgenes downregulate expression of GLR glia reporters for *nep-2*/neprilysin, *hlh-1*/MyoD/Myf, *snf-11*/*GAT* GABA transporter, *unc-46/*LAMP-like, *gly-18*/N-acetylglucosaminyl transferase, and *lgc-55*/tyramine receptor (**Figs. 3B-G**). Expression of *unc-30/Pitx2*, is not affected by *let-381* RNAi (see below), allowing us to visualize the cells and determine that they are still generated. Indeed, *let-381* RNAi animals neither display an anteriorly-displaced nerve ring nor have extra head muscle cells (**Appendix Figs. S3A and B**). Thus, LET-381 is required post-embryonically to maintain GLR glia gene expression, and this function is distinct from its role in generating GLR glia.

**Figure 3.**
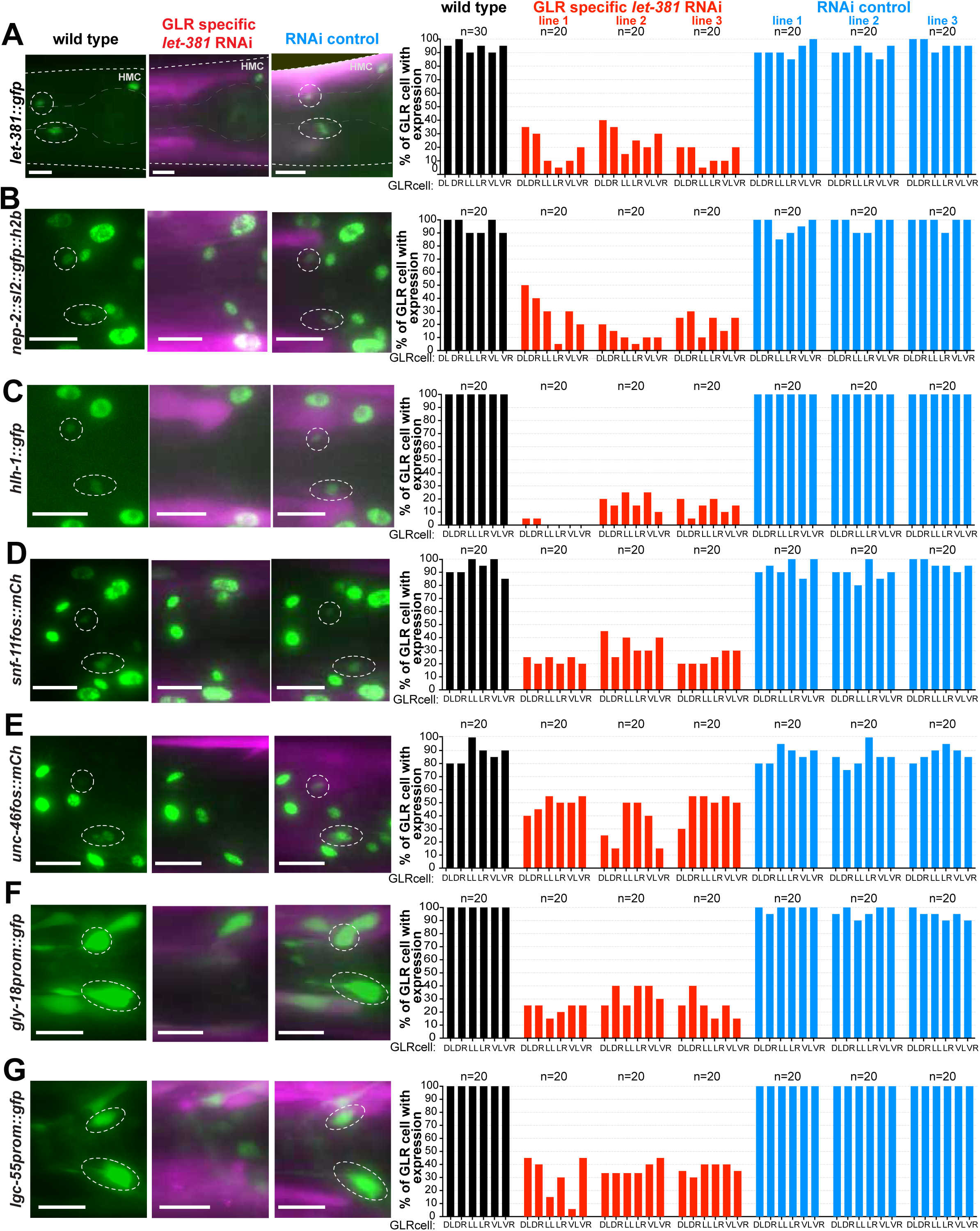
Postembryonic *let-381* knockdown affects GLR glia gene expression. **(A – G)** *gfp* or *mCherry*-based reporter expression (green) in GLR glia of endogenously tagged (A) *let-381*, (B) *nep-2* and (C) *hlh-1* and transgenic (D) *snf-11* (E) *unc-46*, (F) *gly-18* and (G) *lgc-55* in wild type (left column), GLR-specific postembryonic *let-381* RNAi (middle column) and RNAi control animals (right column). Fluorescence images of L4 animals are shown. GFP expression in GLR glia (dashed white circles) is downregulated in the *let-381* RNAi animals but not affected in RNAi control. RNAi and control lines carry a co-injection marker expressed in body wall muscle (magenta). Quantification is shown in bar graphs on the right. Each bar represents % of expression in each of the 6 GLR glia (DL, DR, LL, LR, VL, VR) in the three different backgrounds (wild type = black, GLR-specific RNAi = red, RNAi control = blue). Three independent extrachromosomal lines were scored for the RNAi and RNAi controls. Data information: Anterior is left, dorsal is up and scale bars are 10 μm for all animal images.

To probe dynamic functions of *let-381*, we used CRISPR/Cas9 to insert sequences encoding an auxin inducible degron (AID) ^36^ into the *let-381* genomic locus (**Fig. 2A**). In the presence of GLR glia-expressed TIR-1 and a synthetic auxin analog (K-NAA), AID-tagged LET-381 protein is rapidly degraded ^37^ specifically in GLR glia (**Fig. 4A**). A three-day exposure to K-NAA starting either at the L1 or late-L4/young-adult stages downregulates *nep-2prom7::rfp* reporter expression (**Figs. 4B-F**). Thus, *let-381* functions cell autonomously and is continuously required to maintain GLR glia gene expression, even in adults.

**Figure 4.**
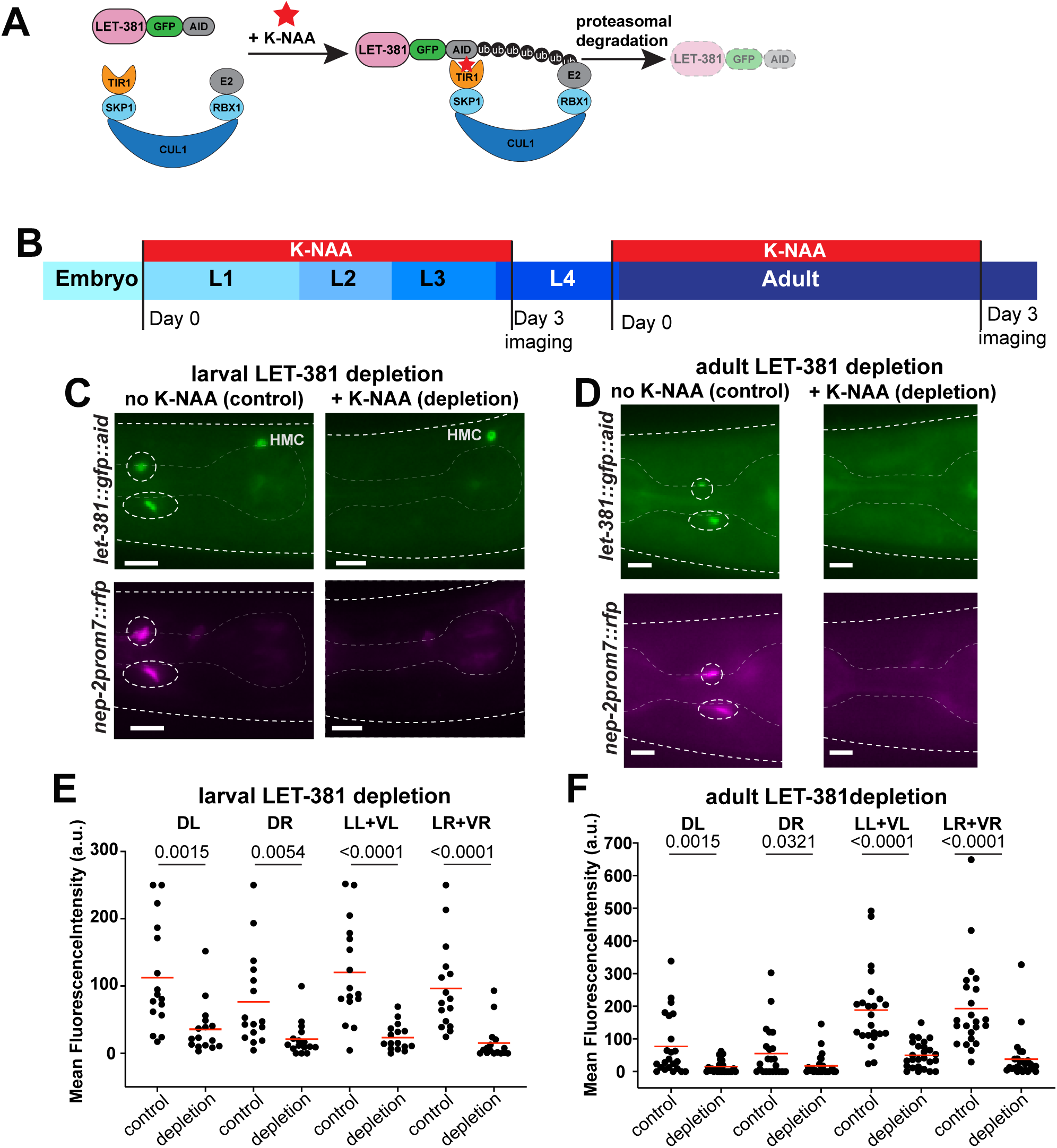
Acute larval and adult LET-381 depletion results in loss of GLR gene expression. (A) Schematic representation of auxin induced degradation ^36^ of LET-381. TIR1 is transgenically provided and expressed specifically in GLR glia by the *nep-2prom7* promoter. (B) Timeline of the auxin inducible depletion experiment. Synchronized populations of L1 or late L4/Young adult animals were placed on plates containing the auxin analog K-NAA and imaged after a three day exposure to K-NAA. Fluorescence intensities of gene expression in GLR glia were compared to age matched animals grown on control plates without K-NAA. (C), (D) Fluorescence images showing the result of (C) larval and (D) adult depletion of LET-381 using auxin inducible degradation. Animals grown on control plates without K-NAA (left panels) show expression of *let-381::gfp* (green) and *nep-2prom7::rfp* (magenta) in GLR glia (dashed white circles). Age matched animals grown on K-NAA containing plates (right panels) show depletion of endogenous *let-381::gfp* expression specifically in the GLR glia; as shown expression in HMC remains unaffected. As a result, expression of the GLR-specific *nep-2prom7::rfp* reporter is downregulated in GLR glia. (E), (F) Quantification (mean fluorescence intensity in cell bodies) of (E) larval and (F) adult LET-381 depletion on expression of *nep-2prom7::rfp* reporter in each GLR glia. Cell bodies of Lateral and Ventral GLR glia are too close to be clearly distinguished, thus they were grouped (LL+VL, LR+VR) for quantification purposes for this experiment. Red lines in dot plots indicate averages. Data information: Un-paired t-test used for statistical analysis in (E) and (F). n=16 for control and depletion in (E), n=23 for control and n=26 for depletion in (F). Anterior is left, dorsal is up and scale bars are 10 μm for all animal images.

### LET-381 binding motifs are required and sufficient for GLR glia gene expression

The expression of LET-381 in GLR glia throughout its life suggests it may function as a terminal selector, directly co-regulating expression of terminal differentiation gene batteries via a shared cis-regulatory motif ^38^. To test this idea, we used sequences identified in our promoter dissection studies, ranging in size from 125 – 1022 bp, as input for the motif discovery tool MEME ^39^. This analysis identified a TGTTTA(C/T/G)A sequence common to all sequence inputs (**Fig. EV1A**). Remarkably, this sequence is highly similar to a previously identified FoxF binding sequence in mice^40^ and to a *C. elegans* LET-381 binding sequence identified through protein binding microarrays ^41^ (**Figs. EV1B and C**). Notably, we find this motif in regulatory regions of all genes whose expression in GLR glia is downregulated by *let-381* knockdown (**Fig. EV1D**). We refer to the TGTTTA(C/T/G)A sequence as the *let-381* motif.

To assess how the motif affects gene expression, we used CRISPR/Cas9 to mutate it in different genomic locations (**Fig. EV1F**). As shown in Fig. 5, mutating *let-381* motifs in upstream regulatory regions of the *nep-2* and *pll-1* genes fused endogenously to *gfp*, significantly reduces *gfp* expression (**Figs. 5A and B**). Animals homozygous for an endogenous *inx-18::gfp* insertion allele we generated localize GFP in bright puncta marking the gap junctions between the GLR glia and the RME motoneurons. Mutagenesis of the *inx-18 let-381* motif eliminates these bright puncta (**Fig. 5C**). Finally, disrupting either of two motifs in the gene *hlh-1* only slightly reduces endogenous *hlh-1::gfp* expression; however, disrupting both together completely abolishes expression (**Fig. 5D**), suggesting that in some contexts, LET-381 binds multiple sites in the same gene.

**Figure 5.**
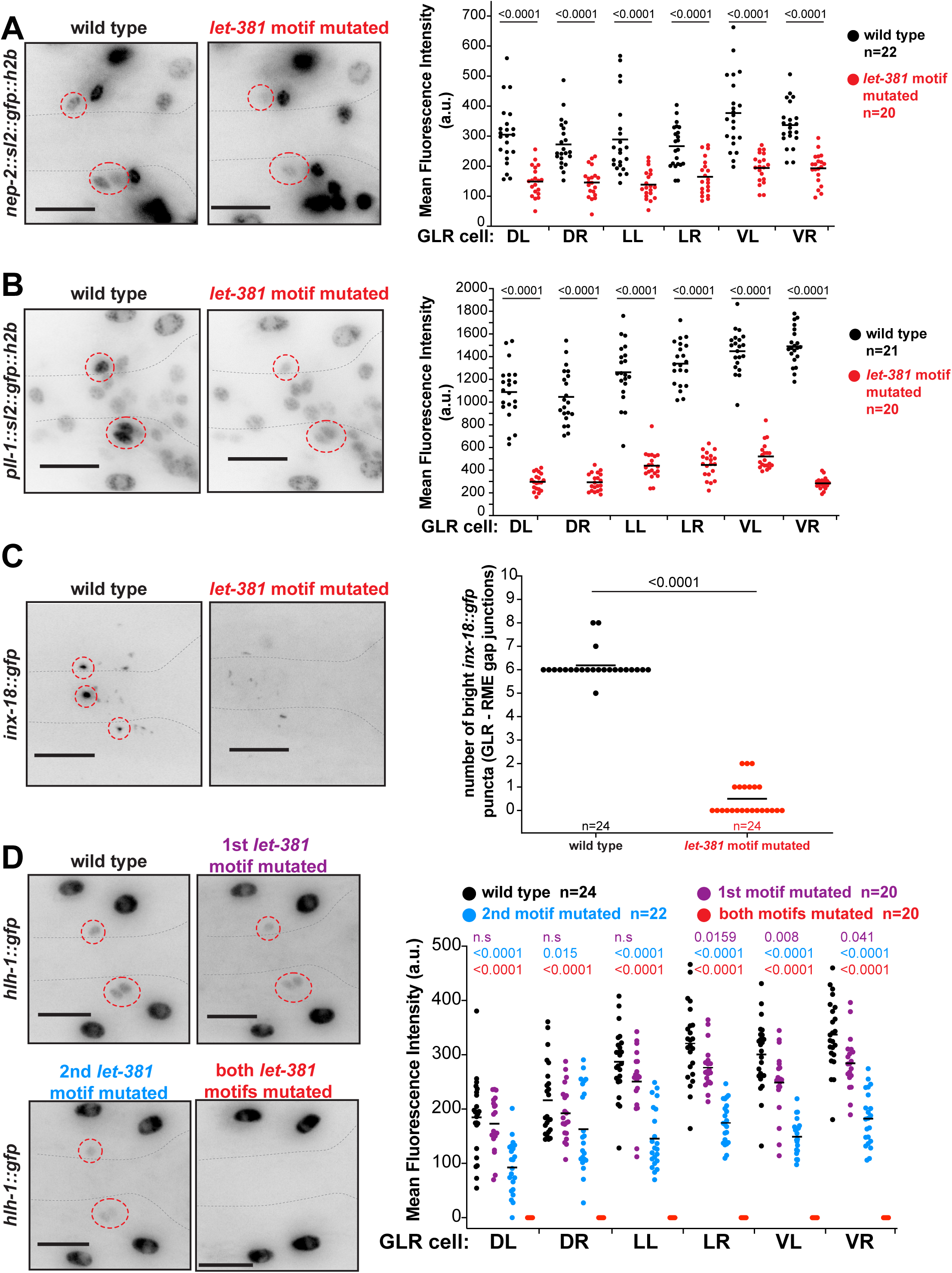
*let-381* motifs are required for endogenous gene expression in GLR glia. (A-D) Endogenous expression of (A) *nep-2*, (B) *pll-1*, (C) *hlh-1* and (D) *inx-18* in wild type and *let-381* motif-mutated animals (details on endogenous gfp reporters and molecular identity of *let-381* motif mutations are shown in Fig. EV1F). Animal images are on the left. Dashed circles outline expression in GLR glia. Quantifications are shown in the dot-plots on the right. Data information: Mean fluorescence expression intensity for each GLR glia cell for (A), (B) and (D) and number of bright gap junction puncta for (C) is compared between the wild type and *let-381*-motif mutated backgrounds. Black lines indicate averages. Un-paired t-test used for statistical analysis. Anterior is left, dorsal is up and scale bars are 10 μm for all animal images.

To further test the idea that *let-381* is a terminal selector gene, we identified motifs in 15 GLR glia-enriched genes whose expression was not previously verified. We then generated animals transgenic for sequences surrounding *let-381* motifs from each gene fused to *gfp* coding sequences (**Table EV1**). We found that *let-381* motif-containing regions (ranging in size from 149 - 254 bp) from 14/19 genes are sufficient to drive GFP expression in GLR glia. Our experiments, therefore, support the idea that *let-381* is a terminal selector gene, controlling coordinate expression of genes expressed in differentiated GLR glia.

### LET-381 regulates its own expression to maintain GLR glia identity

Given the requirement for *let-381* in maintaining GLR glia gene expression even in adults, we wondered how *let-381* expression itself is maintained. We identified a conserved *let-381* motif upstream of the *let-381* first exon (**Figs. EV2A and B**). We wondered whether, through positive feedback, this element could account for sustained *let-381* expression through adulthood, and used CRISPR/Cas9 to delete this motif. In mutant animals, *let-381(ns1026)*, endogenous *let-381::gfp* expression is observed in GLR glia of bean-stage embryos at levels similar to wild type. However, expression gradually wanes and is completely lost in L1 larva and older animals (**Fig. 6A**). Thus, *let-381* is required to maintain its own expression through an autoregulatory *let-381* motif. Expression of nine downstream GLR glia genes (but not *unc-30*, see below) is also gradually lost by the L1 stage (**Figs. 6B-E, and Figs. EV2C-G**), further supporting the notion that LET-381 is required for GLR glia fate maintenance. Of note, *let-381* auto-regulation mutants exhibit neither an anteriorly displaced nerve ring nor extra head muscles, consistent with the RNAi and AID knockdown results.

**Figure 6.**
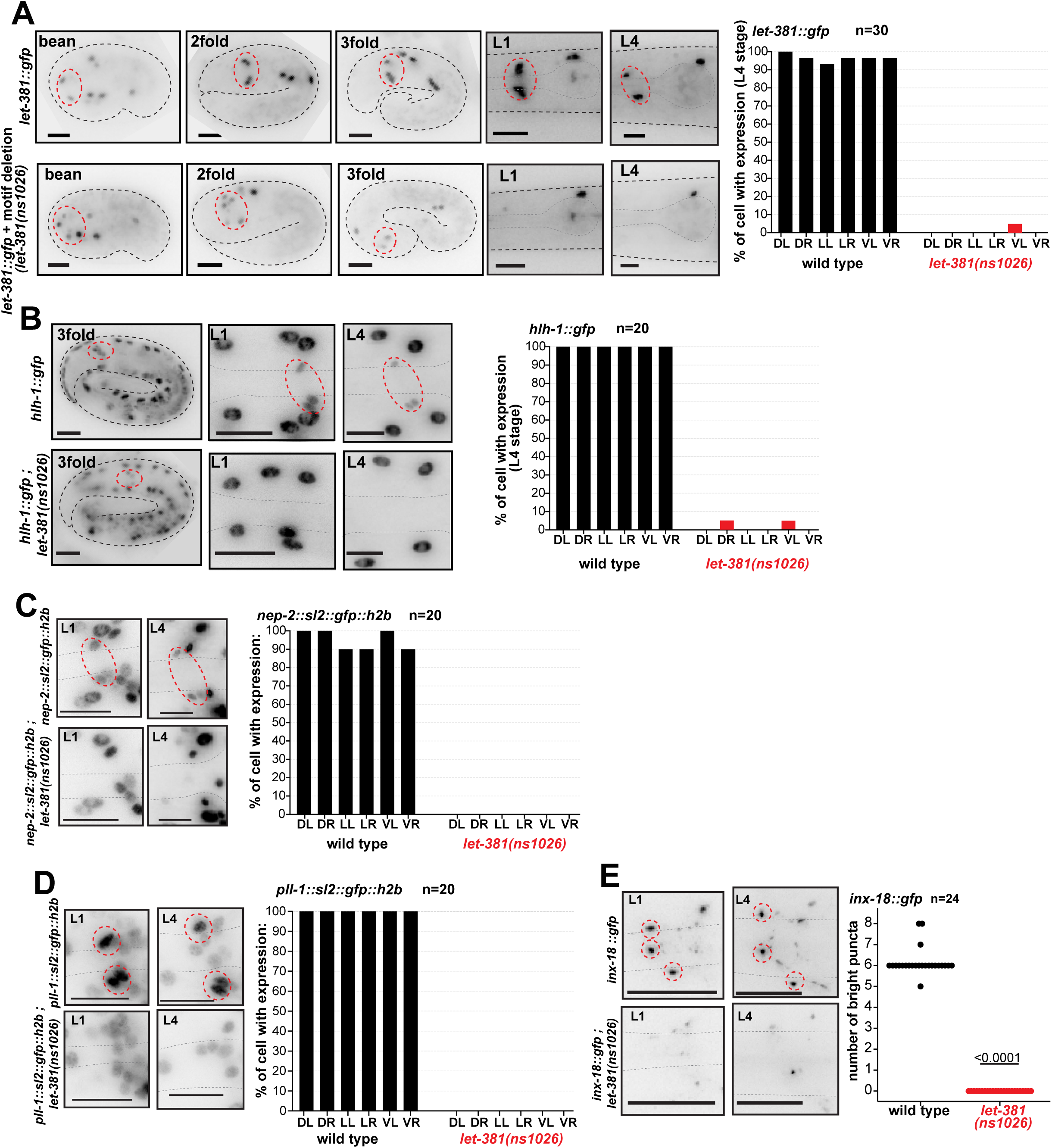
*let-381* positively regulates its own expression. (A) Endogenous *let-381::gfp* expression in different development stages in 2 different backgrounds: wild type (top), *let-381* autoregulatory motif deletion (bottom). Animal images and are shown on the left and quantification of expression in the L4 stage is shown on the right: percentage of each GLR cell expressing *let-381::gfp* in wild type and autoregulatory motif deletion backgrounds. (B) – (E) Effect of autoregulatory motif deletion on endogenous (B) *hlh-1*, (C) *nep-2*, (D) *pll-1* and (E) *inx-18 gfp* reporter expression in GLR glia at different developmental stages. Quantification is shown on the right of each animal image panel for L4 animals. Red dashed circles outline GLR glia. Data information: Un-paired t-test used for statistical analysis in (E). Anterior is left, dorsal is up and scale bars are 10 μm for all animal images.

### *unc-30/Pitx2,* a *let-381/FoxF* target, controls GLR glia gene expression and shape

Our transcriptome studies revealed that transcripts encoding the transcription factor UNC-30/Pitx2 are also significantly enriched in GLR glia. UNC-30 was previously identified as a terminal selector of GABAergic identity in ventral cord neurons ^42–44^, and a recent study showed that GLR glia are GABA positive by immunostaining and express the GABA related genes *snf-11/GAT*, *gta-1/GABA-T* and *unc-46/LAMP*^17^. We found that animals carrying an endogenous *unc-30::gfp* reporter we generated (**Fig. 7A**), express GFP in GLR glia starting at the embryonic bean-stage and through to adulthood (**Fig. 7B**).

**Figure 7.**
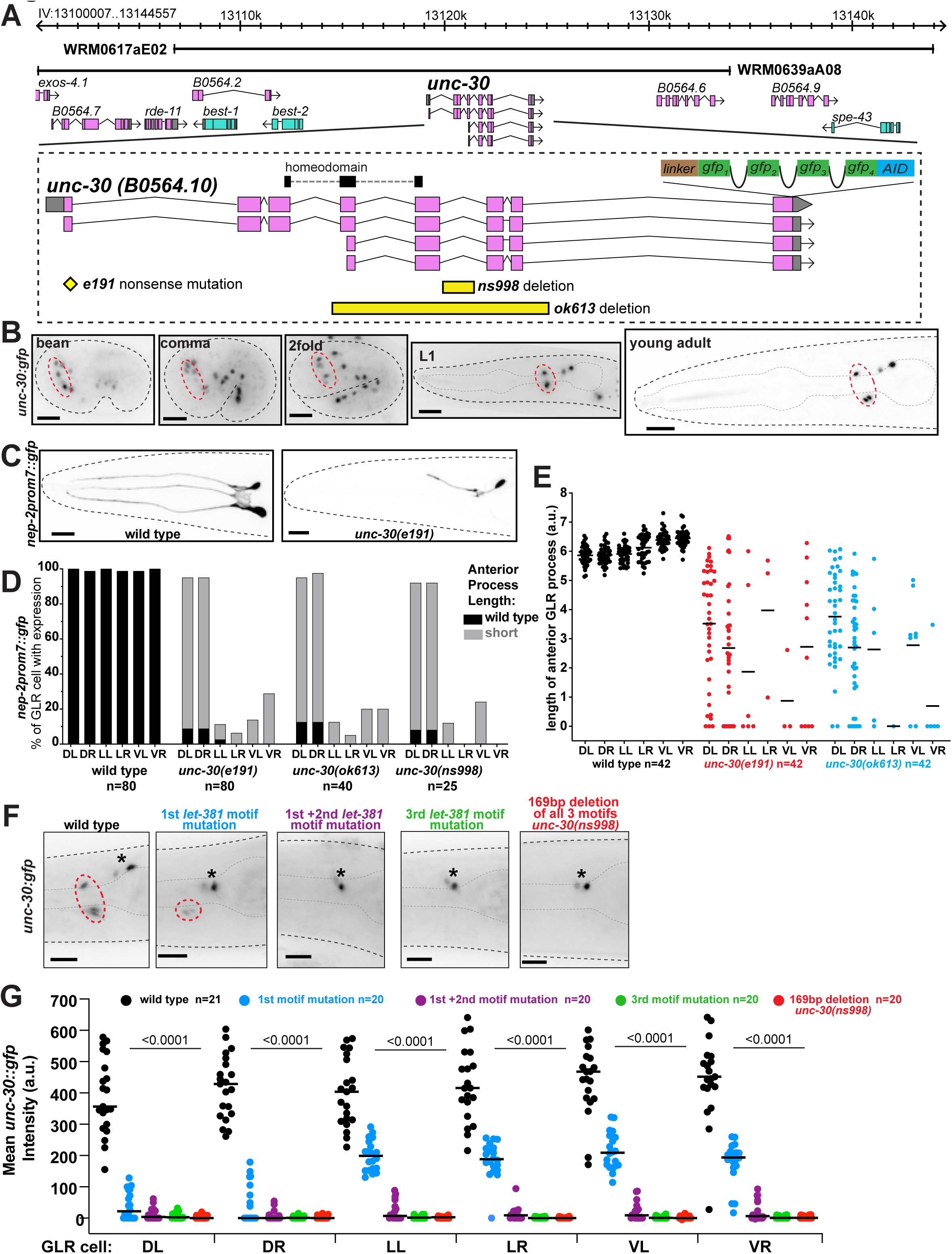
*unc-30* acts downstream of *let-381* to control GLR glia gene expression and the length of GLR anterior process. (A) *unc-30* genomic locus showing mutant alleles, reporters, fosmid genomic clones used in this study. (B) Expression of the endogenous *unc-30::gfp* reporter at different stages during development. Dashed red circles outline GLR glia. (C) *nep-2prom7::gfp* expression in a wild type L4 (left) and a *unc-30(e191)* null mutant L4 animal (right). GFP expression is lost in the lateral and ventral GLR glia in *unc-30(e191)*. The anterior process of the dorsal GLR glia still expressing GFP is shorter than that of a wild type animal. (D) Quantification of percentage of each GLR glia cell with *nep-2prom7::gfp* expression (L4 stage) for different unc-30 mutant backgrounds. (E) Quantification of the length of GLR anterior processes (L4 stage) for different *unc-30* mutant backgrounds. (F) Images of L4 animals showing differences in endogenous *unc-30::gfp* expression upon mutation of the *let-381* motifs present in the fifth intron of *unc-30*. Dashed red circles outline GLR glia and black asterisks denote expression in neurons. (G) Quantification of *unc-30::gfp* fluorescence intensity for each GLR glia cell; black lines in bar graphs indicate averages. Data information: Un-paired t-test used for statistical analysis in (G). Anterior is left, dorsal is up and scale bars are 10 μm for all animal images.

To determine whether *unc-30* controls GLR glia gene expression, we crossed gene reporters into *unc-30(e191)* mutants, harboring an early nonsense mutation (**Fig. 7A**). Endogenous *hlh-1::gfp* expression is completely abolished in these animals, and expression of endogenous *nep-2::gfp*, transgenic *nep-2prom7::gfp*, or *lgc-55prom::gfp* is also lost, but mainly in lateral and ventral GLR glia (**Figs. 7C and D, and Figs. EV3B-D**). Expression of transgenic *gly-18prom::gfp*, *pll-1prom1::gfp* and *snf-11f^osmid^::SL2::mCherry:H2B* reporters is reduced but not abolished (**Fig. EV3E-G**). In most animals in which reporter expression is not extinguished, GLR glia anterior processes are shortened (**Figs. 7C and E**). Similar findings are observed with the *unc-30(ok613)* deletion mutant (**Figs. 7A, C and E**), and all GLR glia defects are rescued with transgenes carrying the wild-type *unc-30* locus (**Fig. EV3A**). Thus, UNC-30 regulates GLR glia gene expression to a more limited extent than LET-381 and also controls GLR glia morphology.

How might LET-381 and UNC-30 interact? We found that the fifth intron of *unc-30* contains three conserved *let-381* motifs located within an 88 bp sequence (**Fig. EV3H and I**). Sequences derived from this intron are sufficient to promote GFP expression in GLR glia (**Fig. EV3H**). Furthermore, mutating the upstream *let-381* motif alone substantially reduces endogenous *unc-30::gfp* expression, and simultaneous mutation of the two upstream motifs, a mutation of the downstream motif alone, or a 169 bp deletion containing all three motifs all specifically abolish *unc-30::gfp* expression in GLR glia, but not in other *unc-30* expressing cells (**Figs. 7F and G, and Fig. EV3I**). Remarkably, the 169 bp deletion allele, *unc-30(ns998)*, phenocopies the effect of *unc-30* null alleles on GLR glia gene expression and anterior process length (**Fig. 7D, and Fig. EV3B**). We found no effects of *unc-30* mutants on the expression of *let-381*; however, *let-381* expressing cells are often displaced along the dorsoventral and left-right axes in these mutants (**Figs. 8A and B**). Together, these results support the conclusion that *unc-30* acts cell-autonomously to control GLR glia gene expression and anterior process length and that *let-381* controls *unc-30* expression in GLR glia. Previous studies suggest that UNC-30 may regulate its own expression^38,44^, suggesting that following initial LET-381 binding, *unc-30* may become independent of *let-381*. This notion is supported by our findings that *unc-30::gfp* expression in GLR glia is unaffected when *let-381* is knocked down post-embryonically (**Fig. EV3J and K**).

**Figure 8.**
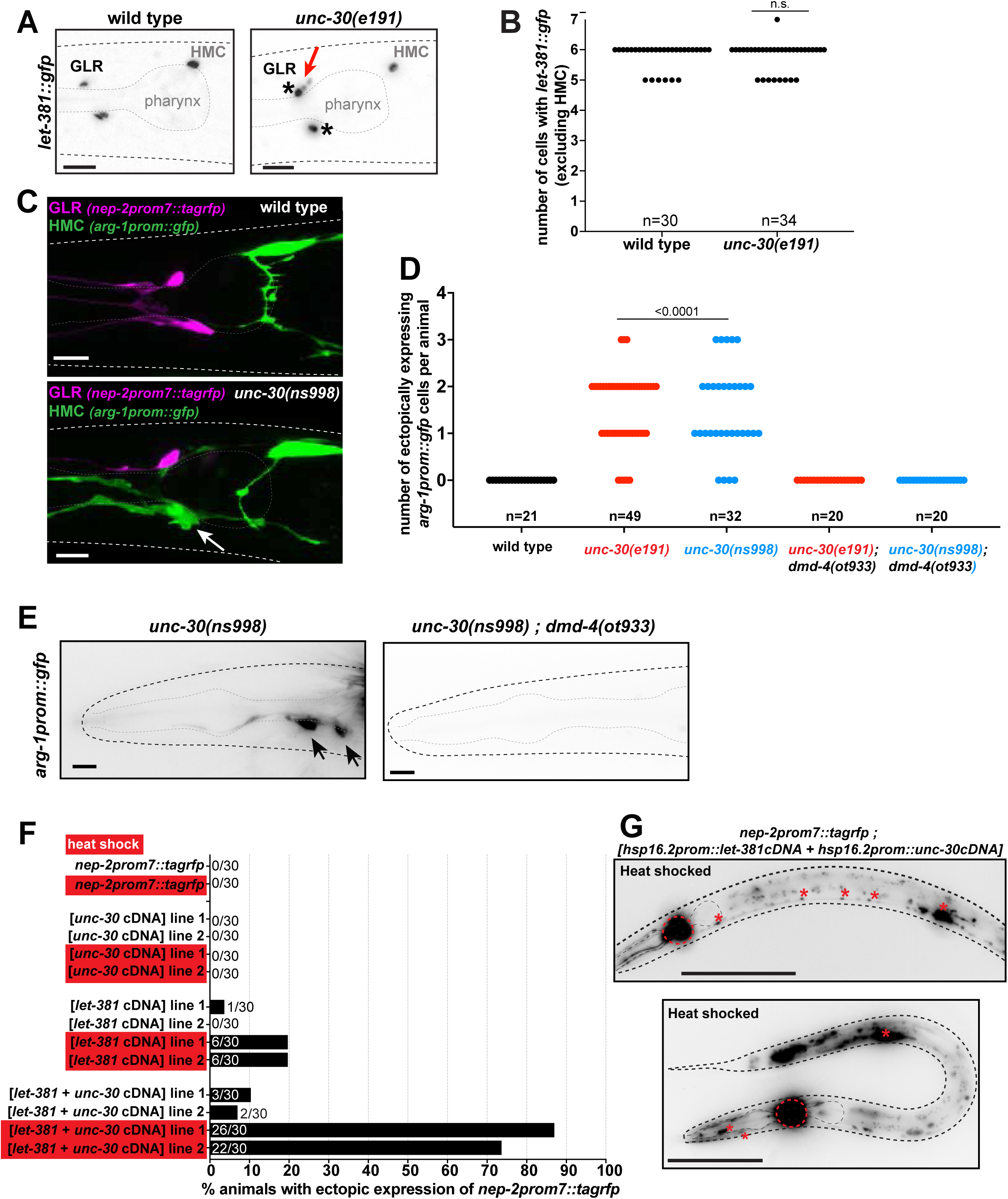
*unc-30* represses HMC gene expression in GLR glia. (A) Images showing *let-381::gfp* expression in wild type (left) and *unc-30(e191)* mutants (right). Expression is observed in GLR glia anterior to the pharynx bulb and the HMC above and posterior to the pharynx bulb. In wild type background animals, GLR glia have a small sesame-like nucleus shape. In contrast in *unc-30* mutants, some GLR glia nuclei appear larger and more round (black asterisks), reminiscent to the nucleus of the HMC cell. In addition, GLR glia are often mispositioned along the dorsoventral or left-right axis in *unc-30* mutants. Red arrow points to dorsally mispositioned cells. (B) Number of *let-381::gfp* expressing cells is unaffected in *unc-30(e191)* null mutants. (C) Expression of GLR-specific *nep-2prom7::tagrfp* (magenta) and HMC-specific *arg-1prom::gfp* (green) in wild type and *unc-30(ns998)* mutant backgrounds. Fluorescence images of L4 animals are shown. In the GLR-specific *unc-30(ns998)* mutant background, GLR glia lose GLR-specific RFP expression and ectopically express HCM-specific GFP instead (white arrow). (D) Quantification of ectopic expression of the HMC-specific *arg-1prom::gfp* in GLR glia in *unc-30(e191)* and *unc-30(ns998)* mutants. (E) Ectopic *arg-1prom::gfp* expression in *unc-30* mutants is not observed in the *dmd-4(ot933)* mutant background. Black arrows point to ectopic *arg-1prom::gfp* expression. (F) Percentage of animals displaying ectopic expression of the GLR glia-specific reporter *nep-2prom7::rfp*, upon heat shock induced misexpression of *let-381* and *unc-30*. Heat shocked animals (red boxes) are compared to age matched non heat shocked controls. (G) Images showing animals with ectopic *nep-2prom7::rfp* expression after heath shock induced misexpression of LET-381 and UNC-30. Red dashed circles outline expression GLR glia. Red asterisks point to expression in ventral nerve cord motor neurons and stomatointestinal muscle (upper panel) and body wall and pharynx muscle (bottom panel). Data information: Un-paired t-test used for statistical analysis in (B) and (D). Anterior is left, dorsal is up and scale bars are 10 μm for (A), (C) and (E). Scale bars are 100 μm for (G).

### *unc-30/Pitx2* represses expression of HMC genes in GLR glia

Wild type GLR glia have small, sesame-seed shaped nuclei. We noticed that in *unc-30* mutants, GLR glia nuclei are larger and rounder, resembling the nucleus of the head mesodermal cell (HMC), another *let-381*-expressing cell (**Fig. 8A**). To determine whether *unc-30* loss results in GLR glia acquiring additional HMC characteristics, we generated a *nep-2prom7::tagrfp* GLR glia reporter strain also expressing *arg-1prom::gfp*, an HMC reporter. We found that in *unc-30(e191)* mutants, some GLR glia that lose RFP expression now ectopically express the HMC reporter. *unc-30(ns998)* mutants exhibit similar defects (**Figs. 8C and D**). Likewise, two additional HMC reporters, *glb-26prom::gfp* and *dmd-4::his24::mCherry*, are also mis-expressed in GLR glia of *unc-30* mutants (**Figs. EV4A-C**). While HMC-converted GLR glia retain *let-381::gfp* expression (**Fig. EV4D**), we never observe GLR glia expressing a mix of GLR glia-specific and HMC-specific reporters. Finally, we found that ectopic expression of the HMC reporter *arg-1prom::gfp* requires the DMRT transcription factor DMD-4 (**Figs. 8D and E, and Fig. EV4E**), normally required for *arg-1prom::gfp* expression in the HMC ^45^. We conclude that UNC-30 acts in GLR glia to repress *dmd-4*-dependent HMC-specific gene expression. This model succinctly explains the differential effect of *unc-30* loss on the various reporters we tested (**Figs. EV4B-G**): *snf-11*, *gly-18* and *pll-1* reporters are normally expressed in both GLR glia and HMC and are therefore less affected by an *unc-30* mutation than *nep-2*, *lgc-55*, or *hlh-1*, which are expressed in GLR glia but not in HMC.

Does *unc-30* control of GLR glia-specific gene expression require inhibition of HMC gene expression? We found that even though the total number of cells expressing *let-381::gfp* is unaltered between wild type and *unc-30* mutants (6 total cells – excluding HMC; **Fig. 8B**), the number of cells expressing either a GLR glia or an ectopic HMC reporter rarely adds up to 6 (**Fig. EV4F**), indicating that there are GLR glia that lose GLR glia gene expression without acquiring HMC fate. Thus, it is likely that *unc-30* mediates GLR glia-specific gene expression independently of HMC gene expression suppression.

### *let-381* and *unc-30* are together sufficient to induce GLR glia gene expression

The broader effect of *let-381* loss on GLR glia gene expression suggests that expression of this gene in naïve cells should drive the GLR glia expression program in these cells. Surprisingly, we found that this is not the case (**Fig. 8F**). Indeed, broad inducible misexpression of *let-381* cDNA using a heat-shock promoter results in minimal misexpression of the *nep-2prom7::tagrfp* GLR glia reporter. We found a similar result using *unc-30* cDNA alone. However, misexpression of both *let-381* and *unc-30* results in highly penetrant misexpression of *nep-2prom7::tagrfp* in many cells including body wall muscle, pharyngeal muscle, the stomato-intestinal muscle and ventral nerve cord motoneurons (**Figs. 8F and G**). Thus, *let-381* is not sufficient to induce GLR glia fate and must cooperate with *unc-30*.

### GLR glia defective animals display locomotion abnormalities

Our development of tools to manipulate gene expression and function specifically in GLR glia allowed us to interrogate the functions of these cells. To do so, we generated a *nep-2prom7*::*egl-1* transgenic line driving postembryonic expression of EGL-1, a caspase activator, only in GLR glia. We observed GLR glia loss, rescued by the *ced-3(n171)* caspase mutation, confirming that GLR glia in this strain die by apoptosis (**Figs. 9A-C**). We next recorded movies and analyzed locomotion of GLR glia-ablated animals freely moving on an agar plate without food ^46,47^. Remarkably, we found that while GLR glia-ablated animals can move, they exhibit severe defects in all locomotion parameters we analyzed, including reduced locomotion rates, increased turning frequency, increased reversal rates, and increased pausing (**Figs. 9D-K, and Movies EV1 and EV2**). These defects are similar to those observed following ablation of CEPsh glia, *C. elegans* astrocytes that line the outer aspect of the nerve ring and which also regulate synaptic function ^46,47^. In addition, while wild type animals exposed to high salt concentrations are initially paralyzed and then recover, GLR glia ablated animals paralyze more quickly and recover at a much slower rate (**Fig. 9L**), suggesting that GLR glia may play an important role in the control of solute permeability and ionic balance in the nerve ring. Thus, GLR glia are important for coordinated locomotion and nerve ring function.

**Figure 9.**
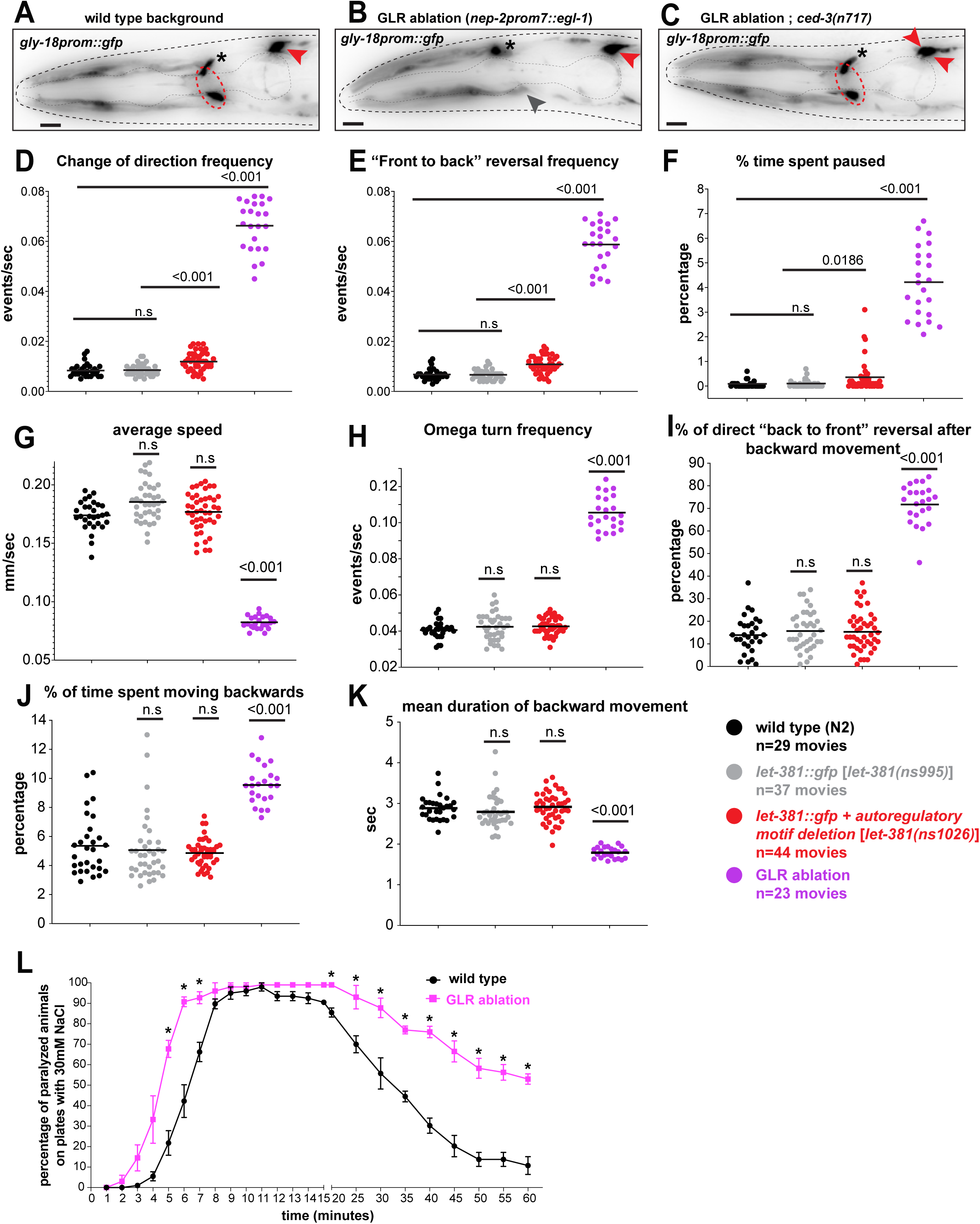
GLR glia-defective animals display locomotion abnormalities and hypersensitivity at high salt concentrations. (A) – (C) Images of L4 animals showing expression of *gly-18prom::gfp* reporter in (A) wild type, (B) GLR glia-ablation and (C) GLR glia-ablation ; *ced-3(n717)* backgrounds. GLR glia are outlined by red dashed circles. Black asterisks denote expression in a neuronal cell just above the dorsal GLR glia and red arrowheads point to HMC cells. In the GLR glia-ablation background the nerve ring is anteriorly displaced as noted by a head muscle arm (grey arrowhead) penetrating the nerve ring at the anterior pharynx bulb. The HMC sister cell that normally dies by programmed apoptotic cell death in a wild type animal, survives and also expresses *gly-18prom::gfp* in the *ced-3(n7171)* mutant background in (C). (D) – (K) Locomotion parameters of foraging wild type (black), control (grey), GLR glia-defective (red) and GLR glia-ablated (purple) animals were analyzed using automated tracking ^46,47^. (D) change of direction frequency, (E) “front to back” reversal frequency, (F) % time spent paused, (G) average speed, (H) omega turn frequency, (I) % direct “back to front” reversal after backward movement, (J) % time spent moving backwards, (K) mean duration of backward movement. (L) GLR glia-ablated animals paralyze at a significantly higher rate than wild type animals when exposed to 300mM NaCl. GLR glia-ablated animals also recover motility at a significantly lower rate. Data information: Un-paired t-test was used for statistical analysis in (D) – (K), * < 0.05 in (F). Anterior is left, dorsal is up and scale bars are 10 μm for all animal images.

We noted that, like animals whose GLR glia progenitors are laser ablated during embryogenesis ^34^, *nep-2prom7*::*egl-1* ablated animals have anteriorly-displaced and sometimes defasciculated nerve rings (**Fig. EV5A**). To determine whether the abnormal locomotion we observed is a consequence of these structural defects, we examined the behavior of axon-guidance mutant strains that also possess anteriorly-displaced nerve rings ^48,49^. Indeed, these mutants also exhibit defects in all locomotion parameters tested. However, the magnitude of these defects is not always the same as in GLR glia-ablated animals (**Fig. EV5B**). Thus, while some of the locomotion defects may be attributed to structural defects in the nerve ring, it appears that animals lacking GLR glia may be additionally compromised. To uncouple the physical positioning of the nerve ring from other GLR glia functions in *C. elegans* locomotion, we examined movement of animals carrying the *let-381* autoregulatory motif mutation, which does not result in nerve ring displacement. This strain, *let-381(ns1026)*, was generated from the *let-381(ns995)* strain, containing an insertion of *gfp* into the *let-381* locus. As in GLR glia-ablated animals, we find specific defects in *let-381(ns1026)* mutants: animals change direction more frequently than *let-381(ns995)* controls (**Fig. 9D**), display a higher frequency of reversals (**Fig. 9E**), and tend to pause more often (**Fig. 9F**). Other behaviors defective in ablated animals are unaltered (**Figs. 9G-K).** Taken together, our observations support the conclusion that GLR glia regulate *C. elegans* locomotory behavior not only by ensuring nerve-ring positioning, but also in non-structural ways, revealing a previously uncharacterized function for these cells.

## DISCUSSION

We describe here a gene regulatory network for the specification and maintenance of *C. elegans* GLR glia (**Fig. 10A**). Early in development, *let-381*/*FoxF* is required to specify GLR glia fate and to suppress sister-lineage body-wall-muscle gene expression. It does so, in part, by promoting the expression of dozens of GLR glia-enriched genes, all of which possess cis-regulatory LET-381 binding motifs. Such a motif upstream of *let-381* ensures that once turned on, the gene remains continuously expressed, as are its targets. Among these targets is *unc-30/Pitx*, required for expression of some *let-381*-dependent genes and for generating GLR glia anterior processes of appropriate length. *unc-30* also independently prevents expression in GLR glia of genes and traits of the HMC, a non-contractile MS-lineage derived cell ^50^. Together, LET-381 and UNC-30 can bestow GLR glia gene expression onto naïve cells that do not normally express these transcription regulators.

**Figure 10.**
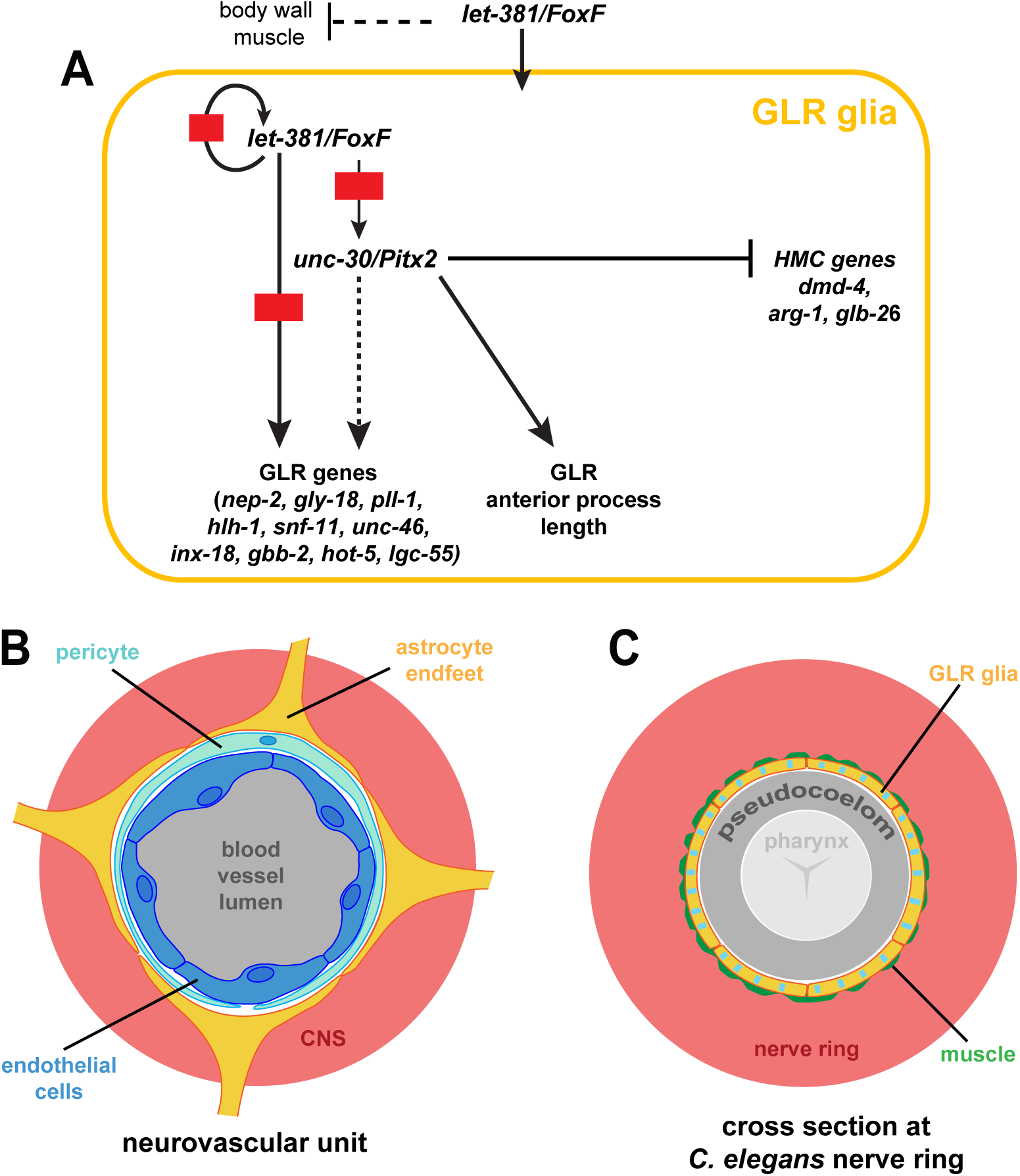
Regulatory network for the specification and identity maintenance of the *C. elegans* GLR glia. (A) Schematic of the regulatory network identified in this study controlling fate specification and differentiation of GLR glia. (B) At the neurovascular unit, endothelial cells (dark blue), pericytes (light blue) and astrocytic endfeet (yellow) form the Blood Brain Barrier isolating the central nervous system (CNS – red) from blood circulation (grey). (C) Similarly, the GLR glia sheet-like processes (yellow with blue stripes to show a mixed astrocytic-endothelial/mural fate) isolate the *C. elegans* nerve ring (red) from the pseudocoelom (grey). A thin layer of head muscle arms (green) penetrates the *C. elegans* nerve ring and therefore GLR flat processes are in close proximity to head neuromuscular junctions ^16^. The pseudocoelom is shown larger than its actual volume.

While transcriptional control of glial cell fate is generally not well understood, neuronal fate specification by terminal selector transcription factors has been studied extensively in *C. elegans*, *Drosophila*, chordates, and mice ^51^. These master regulators, often acting with a specific cofactor ^38,51^, establish and maintain neuron identity by co-regulating a wide variety of neuronal features, including gene expression and synaptic connectivity. Terminal selectors also ensure neuron identity by repressing alternative neuronal fates ^52,53^. Here we show that similar fate control is exercised in *C. elegans* GLR glia by *let-381/FoxF* and its cofactor *unc-30/Pitx2*. It is therefore plausible that such mechanisms also specify the fates of glial cells in other settings, and may account for regional differences between glia in the vertebrate brain.

The formation of intestinal muscle from visceral mesoderm using FoxF transcription factors is conserved among *Drosophila* ^54^, planaria ^32^, *Xenopus* ^55^ and the mouse ^56,57^. In *C. elegans*, *let-381/FoxF* is not expressed in the 4 intestine associated muscles. Nonetheless, it is expressed in GLR glia, the head mesodermal cell (HMC), and the phagocytic coelomocytes, all derived from the mesoderm-like lineage of the blast cell MS. Similar to our findings in GLR glia, *let-381/FoxF* may function with *dmd-4/Dmrt* in HMC (N.S. and S.S., in preparation) and with *ceh-34/Six2* in coelomocytes ^31^. Intriguingly, in planaria, *FoxF1* promotes specification of phagocytic cells, including phagocytic glia ^32^. These findings suggest that FoxF transcription factors are key cell-fate regulators of non-contractile mesodermally-derived cells, including mesodermal glia. Indeed, single-cell sequencing of mouse CNS cells reveals *FoxF1*/*2* expression in blood vessel endothelial and mural cells ^28,58–60^, with *FoxF2* driving pericyte differentiation and maintenance of the blood-brain barrier^33^. While *FoxF* genes are expressed only at low levels in microglia ^59^, it remains possible that these genes direct early microglia differentiation.

Intriguingly, cerebellar astrocytes, which often ensheath blood vessels, also express *FoxF1* ^61^, raising the possibility that *FoxF* genes play a broader role in specifying the entire neurovascular unit. Supporting this idea, we show here that *C. elegans* GLR glia merge astrocyte and mural cell molecular and anatomical characteristics (**Fig. 10, B and C**). In phylogenetically older species, such as sturgeons (subclass: *Chondrostei*), astrocytes are thought to be the main components of the blood-brain barrier ^62^. Might GLR glia, by analogy then, form a barrier that isolates the *C. elegans* nerve ring from the pseudocoelom? Unlike endothelial cells, GLR glia do not adhere to each other to form a tight seal, and gaps between the cells are evident on electron micrographs ^16^. Furthermore, neuronal cell bodies, which reside outside the nerve ring, are not ensheathed and could have direct access to materials within the pseudocoelom. Nonetheless, roles for GLR glia in blocking diffusion of synaptically-released factors into the pseudocoelom, which can then have systemic effects, are plausible, as are more general roles in regulating the extracellular environment of the nerve ring. Indeed, our behavior studies of GLR glia-ablated animals and GLR glia mutants reveal disruptions in the coordination of locomotion, suggesting effects on neuronal signaling within the nerve ring. The similarities in locomotory defects between CEPsh astrocyte-ablated animals and GLR glia-ablated animals suggest that both glial types affect a similar set of underlying processes. CEPsh glia wrap around the outside aspect of the nerve ring, are not in direct contact with the pseudocoelom, and are also not enriched for endothelial genes, further supporting the idea that GLR glia have merged astrocyte and endothelial functions.

## MATERIALS AND METHODS

### Caenorhabditis elegans strains and handling

Animals were grown on nematode growth media (NGM) plates seeded with *E. coli* (OP50) bacteria as a food source unless otherwise mentioned. Strains were maintained by standard methods ^63^. Wild type is strain N2, *C. elegans* variety Bristol RRID:WB-STRAIN:WBStrain00000001. A complete list of strains generated and used in this study is listed below. A few of the strains were previously published, and/or obtained from the Caenorhabditis Genetic Center (CGC), or the TransgeneOme project.

### CRISPR/Cas9 genome editing

CRISPR/Cas9 genome editing was performed using Cas9, tracrRNAs and crRNAs from IDT as described in ^64^.

Generation of deletion alleles was performed by use of two crRNAs and a ssODN donor as follows:

*- unc-30(ns998[*ns959])* [deletion of all three *let-381* motifs from the 5^th^ intron of *unc-30*]:

crRNAs (tctcgtgtggtataaacaat, actcggggtacagataacta) ssODN (gtcaggtaagcagaaggcaggcatcaggagttaattgggaacataattaagaatgaaaaaatatatcaaca)

*- let-381(ns1026[*ns995])* [deletion of *let-381* autoregulatory motif]: crRNAs (tggttgaagagacatacatc, ttatggatggaaaacagacg) ssODN (tcatcatacttttccctctatcttctcaaccagatctgttttccatccataagccaccaccccattctgc). CRISPR/Cas9 generated indel: deletion from –413 to –307 and random insertion of a 34 bp sequence cttatcttctcaatcttctcaaccagatgtgttg.

*- hlh-1(ns1015[*syb3025])* [mutation of the 1^st^ *let-381* motif at +726 from tgtttaca to ccgcgg and a deletion from +743 to +807 containing the 2^nd^ *let-381* motif at +765)

crRNAs (gtgtttacattgtgcaaact, tcttgaaaaattcgtagact) ssODN (atgggaatagtaaagggaggggggtgccgcggttgtgcaaactgggttaacccgttgtaaacataaatcgctaatagga a)

*let-381* motif substitutions were performed by a single crRNA and a ssODN donor containing the desired mutations.

*- nep-2(ns1012[*syb4689])* [substitution of *let-381* motif at –1133 tgtttaca to ccgcgg]

crRNA (caattgaggaacactgggcg) ssODN (cattccgattcccacttggcactgtgccaagttgcgcccagtgttcctcaattgccgcggacagcggctccggggggc)

*- pll-1(ns1040[*syb5792])* [substitution of *let-381* motif at –100 from tctaaata to ccgcggta]

crRNA (acattttggcgtcgacggcg) ssODN (tttctagtagtagcaacagctcacaagacattttgaagcgccgtcgacgccaaaaccgcggtagaaaagaagaaaaaggaaaaaaactggaaacgg)

*- hlh-1(ns1016[*syb3025])* [substitution of the 1^st^ *let-381* motif +726 from tgtttaca to ccgcgg]

crRNA (gtgtttacattgtgcaaact) ssODN (atgggaatagtaaagggaggggggtgccgcggttgtgcaataagccttttaatccattttagtttatttcctttttcttt)

*- hlh-1(ns1027[*syb3025])* [substitution of the 2^nd^ *let-381* motif at +765 from agtttatt to agccatgg]

crRNA (gtgtttacattgtgcaaact) ssODN (aggggagaagagaatttatgaaatgggtcatgggaatagtaaagggaggggggtgtttacattgtgcaaacttttgccttttaatccattttagccatggtcctttttcttttcaattcttgaaaaattcgtagactg)

*- inx-18(ns1010[*syb2879])* [substitution of the *let-381* motif at +364 from tctaaaca to aactgg]

crRNA (ggtcatttctcataggaaga) ssODN (aaacttcttgacatttttggtcatttctcataggaagacttgatttccatggaaacatttttgggcggcggcgggct)

*- unc-30(ns1000[*ns959])* [substitution of the 1^st^ *let-381* motif] crRNA (tctcgtgtggtataaacaat) ssODN

(cagtcaggtaagcagaaggcaggcatcaggagaaaattgtttataccacacgagaatctagaacagtgtcagttttctttcgccccttcttg)

*- unc-30(ns999[*ns959])* [substitution of the 1^st^ + 2^nd^ *let-381* motifs] crRNA (tctcgtgtggtataaacaat) ssODN

(cagtcaggtaagcagaaggcaggcatcaggagaaaattggtaccaccacacgagaatctagaacagtgtcagttttctttcgccccttcttg)

*- unc-30(ns1001[*ns959])* [substitution of the 3^rd^ *let-381* motif] crRNA (cctcttatgtcataaacaattgg) ssODN (ctttcgccccttcttgccagacatcatcttgaatcttatgtcccatggattggaggcgggggtactccgcttgtctcaac)

Generation of the *let-381* and *unc-30* knock in reporters was performed by a single crRNA and a dsDNA asymmetric hybrid-donor containing a linker::gfp::aid cassette for C-terminal tagging (see sequence below – amplified from plasmid pNS7) and 120bp flanking arms with homology to target sequence.

*- let-381(ns995[let-381::gfp::aid])* crRNA (gctattccacaagatttat)

*- unc-30(ns959[unc-30::gfp::aid])* crRNA (aagtggtccactgtactgac)

Cassette sequence:

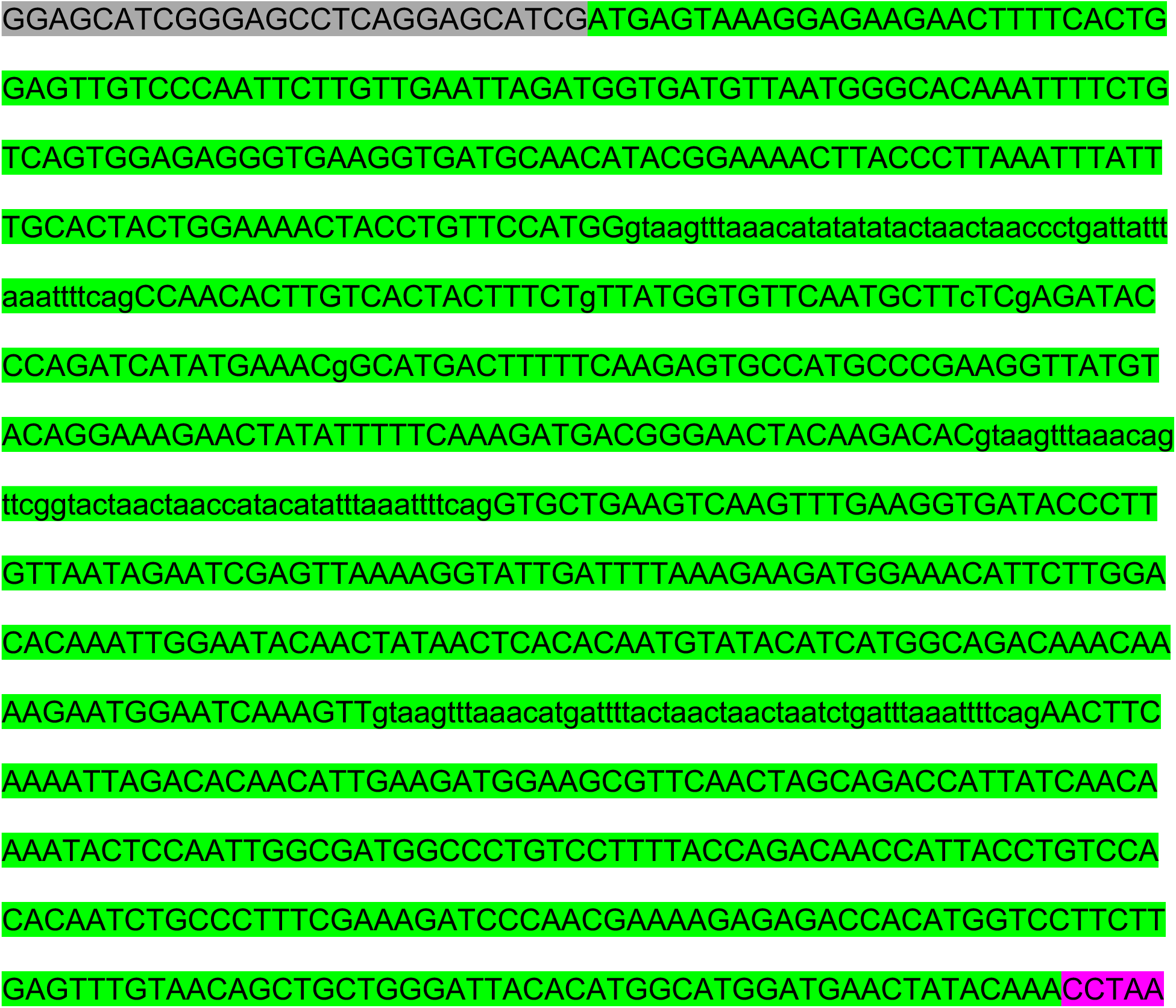
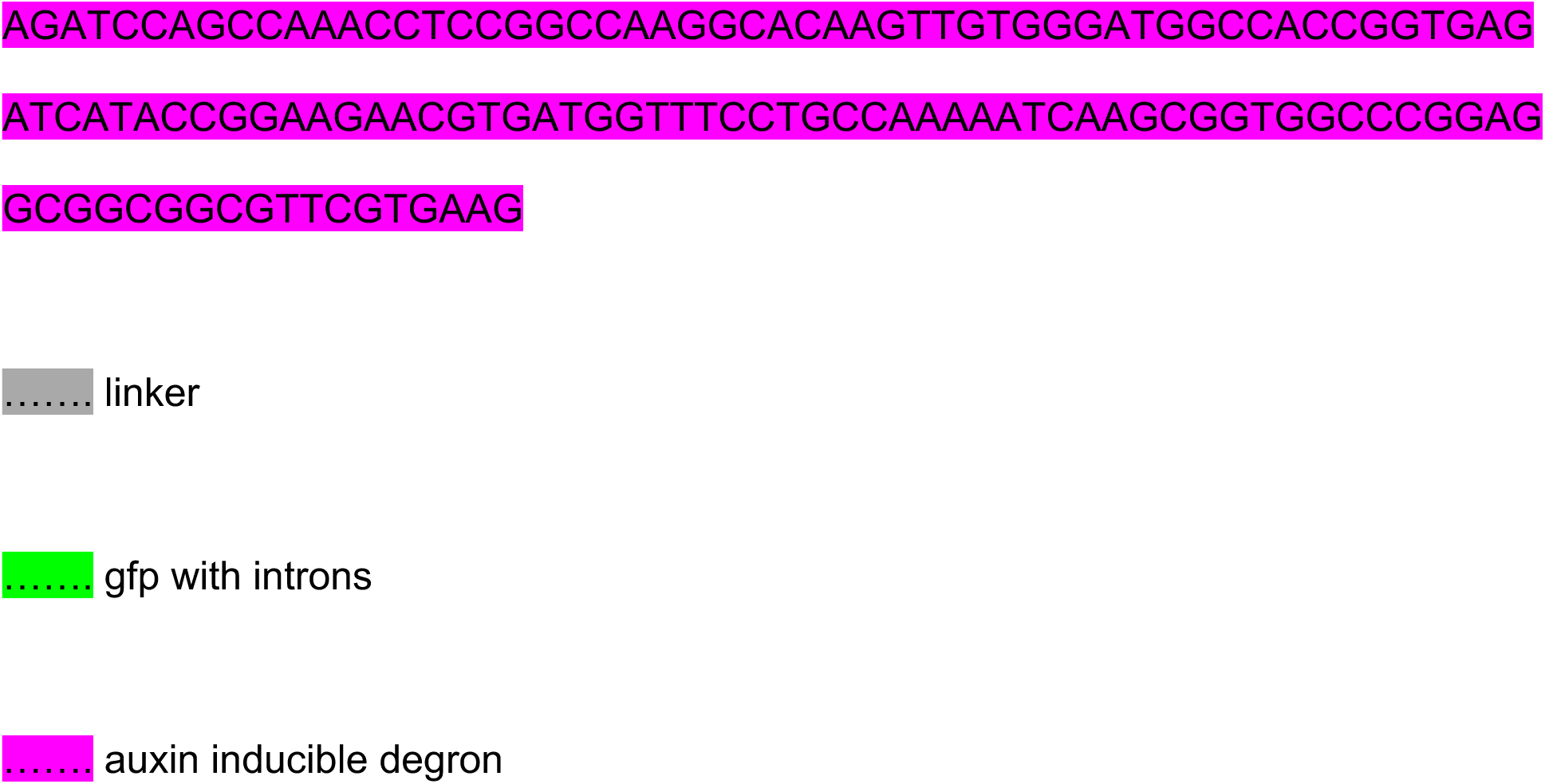

Generation of the *fkh-9* knock in reporter was performed as previously described in ^65^. Cas9, tracrRNA, crRNA and ssODN carrying the 2x(sfGFP_11_) tag were injected, as described in ^64^, into strain CF4587 which ubiquitously expresses sfGFP_(1-10)_. crRNA (ccagtcttttcttcttcaat), ssODN (ctttctccaatcaaggccgagagccagtcttttcttcttcaaggaggaggatccggtgattccggcggcgttgacgtactcgt ggaggaccatgtggtcacgtcctcctcccgtgaccacatggtcctccacgagtacgtcaacgccgccggaatcacctag**a** caccgattctgaactgattgaattagtcgcagttacg)

The following knock in reporter alleles were generated by SUNY Biotech:

*- hlh-1(syb3025[hlh-1::gfp::aid])* C-terminal tagging with *linker::gfp::aid*.

*- inx-18(syb2879[inx-18::aid::emgfp])* tagged with *aid::emgfp* at position +6629bp from *inx-18* start codon.

*- nep-2(syb4689[gfp::h2b::sl2::nep-2])* N-terminal tagging with *gfp::h2b::sl2*.

*- pll-1(syb5792[pll-1::sl2::gfp::h2b])*, *gbb-1(syb5704[gbb-1::sl2::gfp::h2b])*, *gbb-2(syb5759[gbb-2::sl2::gfp::h2b])* C-terminal tagging with *sl2::gfp::h2b*.

### Generation of transgenic reporters and rescuing constructs

All reporter gene fusions for the *cis*-regulatory analysis and the minimal promoters containing *let-381* motifs were generated by a PCR fusion approach ^66^. Genomic promoter fragments were fused to *gfp* or *tagrfp* followed by the *unc-54* 3’ UTR. Promoters were initially amplified with primers A (forward) and B (reverse) and gfp or tagrfp was amplified by primers C (forward) and D (reverse). For the fusion step, amplification was done using primers A* and D* as previously described ^66^. PCR fusion DNA fragments were injected as simple extrachromosomal arrays in a *pha-1(e2123)* mutant background strain in the following concentrations: promoter fusion (50ng/μL), *pha-1* rescuing plasmid (pBX) (50ng/μL). Promoter size and distance from the start codon for each promoter is shown in Fig. 1C for *nep-2*, Appendix Figs. S1A-E for *lgc-55*, *egl-6*, *gly-18, hlh-1*, *inx-18* respectively and Fig. EV3H for *unc-30*. Standard 20bp oligos were used to amplify these promoters from wild type (N2) genomic DNA.

The minimal GLR glia-specific *nep-2prom7* was subcloned by Gibson cloning into the *tir-1* coding plasmid pLZ31 ^36^. *nep-2p7::tir-1::unc-54* 3’UTR was amplified by PCR and injected at 50ng/μL with *unc-122::mCherry* (50ng/μL) as co-injection marker. Similarly *nep-2prom7* was subcloned by Gibson cloning into the *egl-1* expressing plasmid pTB28^67^. *nep-2prom7::egl-1::unc-54 3’UTR* was amplified by PCR and injected at 50ng/μL with *myo-3prom::mCherry* (50ng/μL) as co-injection marker.

Rescuing *let-381* fosmids were injected as simple extrachromosomal arrays at 50ng/μL with a *myo-3prom::mCherry* co-injection marker (25ng/μL) and pBluescript (25ng/μL) as filler DNA to reach a minimum DNA injection concertation of 100ng/μL.

Rescuing *unc-30* fosmids were linearized using NotI (NEB R3189S) and injected as complex extrachromosomal arrays in the following concentrations: linearized fosmid (30 ng/μL), linearized *unc-122prom::gfp* (5 ng/μL) as co-injection marker, sonicated *E.coli* (strain OP50) genomic DNA (100 ng/μL).

Rescuing “*unc-30* whole locus”, including a 2.8kB promoter, *unc-30* gene with introns and 3’UTR, was PCR amplified from genomic DNA and injected as simple extrachromosomal arrays at 50 ng/μL with *unc-122prom::gfp* (40ng/μL) as co-injection marker.

Integrated transgenic constructs (*nsIs700*, *nsIs746*, *nsIs758*, *nsIs831*, *nsIs835*, *nsIs879*, *nsIs854*) were generated by exposure to 33.4 μg/mL trioxsalen (Sigma T6137) and UV irradiation using a Stratagene Stratalinker UV 2400 Crosslinker (360 μJ/cm^2^ x100) as previously described ^68^.

Fosmid reporters were generated as previously described ^69^. More specifically, *gly-18* and *gpx-8* fosmids (WRM0641dF02 and WRM065dE01 respectively) were tagged at the C-terminus of the target genes with SL2::YFP::H2B amplified from pBALU23 ^70^. Fosmid reporters were linearized using NotI and injected in the *pha-1(e2123)* mutant background strain as complex extrachromosomal arrays in the following concentrations: linearized fosmid (30 ng/μL), linearized pBX (pha-1 rescuing plasmid) (2.5 ng/μL), sonicated *E.coli* (strain OP50) genomic DNA 100 ng/μL.

### Temporally controlled, GLR glia-specific LET-381 protein degradation

Exposure of *C. elegans* to a synthetic auxin analog K-NAA, results in ubiquitination and subsequent proteasomal degradation of auxin inducible degron (AID)-tagged proteins ^37^. This occurs only in the presence of transgenically provided TIR-1, the substrate recognition component of the E3 ubiquitin ligase complex. Strains carrying *let-381::gfp::aid* [*let-381(ns995)*] were crossed into a strain expressing *tir-1* specifically in GLR glia (*nsIs879 [nep-2prom7::tir-1]*). K-NAA, 1-napthaleneacetic Acid Potassium Salt (PhytoTech Labs #N610), was dissolved in sterile H_2_O to prepare a 200mM stock solution. OP50-seeded NGM plates were coated with K-NAA for a final concentration of 4mM. Synchronized populations of L1 or late-L4/young-adult animals were transferred on K-NAA plates and grown at 20°C for the duration of the experiment. Age matched animals placed on OP50-seeded NGM plates coated with sterile H_2_O, were used as controls.

### Cell-specific RNAi

A *let-381* fragment, spanning from +421 to +1578 from the *let-381* start codon (Fig. 2A), was amplified by PCR and fused in the sense and antisense orientation under the GLR glia-specific promoter *nep-2prom7* (method described in ^35^). These sense and antisense *let-381* fragments were injected at 40ng/μL each, together with 25 ng/μL of the co-injection marker *myo-3prom::mCherry* driving expression in body wall muscle. Animals injected with unfused *nep-2prom7* (40ng/μL) together with the co-injection marker *myo-3prom::mCherry* (25ng/μL) were used as controls. Three independent extrachromosomal arrays were scored for each genotype. GLR glia are refractory to RNAi; thus, RNAi sensitized background strains carrying *eri-1(mg366)*^71^ were used for these experiments.

### Heat-shock induced misexpression

One step RT-PCR (Invitrogen #12594025) was used to isolate *let-381* and *unc-30* cDNAs (primers for *let-381*: atggaatgctcaacag, ctagcaatccgataaatc, primers for *unc-30*: atggatgacaatacggccac, ctaaagtggtccactgtact), which were subsequently cloned under the heat-shock inducible promoter *hsp-16.2* by Gibson cloning. These constructs were injected at 50ng/μL to generate transgenic lines carrying *hsp-16.2::let-381* cDNA (*nsEx6533*, *nsEx6534*), or *hsp-16.2::unc-30* cDNA (*nsEx6535*, *nsEx6536*) or both (*nsEx6530*, *nsEx6531*) and crossed into a strain expressing RFP specifically in GLR glia (*nsIs700 [nep-2prom7::tagrfp]*). Twenty five 1-Day adult animals (P0) from each genotype were heat shocked by incubating parafilmed plates in a 32°C water bath for 2.5 hours. Heat shocked P0s were kept at 25°C and allowed to lay progeny (F1s) for 15 hours before being removed from the plate. F1 progeny were grown at 25°C for another 24 hours and then scored for ectopic expression of the GLR glia-specific reporter. Age matched non-heat shocked animals were used as controls.

### High salt motility assays

One day adult animals (24 hours after L4 stage, blinded for each genotype) were placed on plates without food, containing 300mM NaCl and scored for their motility over a period of 60 minutes: every one minute for the first 15 minutes and every 5 minutes after that. Animals were score for loss of motility (paralysis) and recovery from paralysis. Twenty to twenty-five animals were tested for each genotype in four replicate experiments.

### Analysis of animal locomotion

Locomotion of wild type, GLR glia-ablated (*nsIs854*), GLR glia-defective *let-381(ns1026*) and control *let-381(ns995)* animals was recorded and analyzed as described in detail in ^46,47^. Briefly, twenty to forty 1-day adult animals were picked to an unseeded plate, washed three times with M9 buffer, and transferred to a 6 cm plate containing 4 ml of NGM-agar without food, with a high-osmolarity barrier (4M Fructose) at the periphery of the plate to prevent wandering of animals off the plate. Animals were allowed to habituate for 20 minutes, then the plate was placed under a camera and locomotion was recorded for 30 minutes at two frames per second and thereafter analyzed as previously described ^46,47^. Several plates were recorded and analyzed for each genotype (29 for wildtype, 37 for control and 44 for GLR glia-defective animals) over the span of three days.

### FACS-based GLR glia cell isolation

For isolation of the GLR glia, synchronized L1 larvae expressing nuclear localized YFP in the GLR glia cells (OS11715) were grown on thirty 10cm plates seeded with HB101 *E. coli* bacteria. After 48 hours at 20°C, L4 larvae animals were washed off plates and subsequently washed ten times with M9 to remove excess bacteria. Each wash consisted of a brief (10 second, 1300 rpm) centrifugation, such that most animals were pelleted, but bacteria stayed in suspension. Animals were then dissociated using SDS-DTT (0.25% SDS; 200 mM DTT; 20 mM HEPES, pH 8.0; 3% sucrose) and Pronase E (15 mg/ml) as previously described^47^. We used 2:1 ratio of SDS-DTT to a volume of packed animals pellet, followed by 4 min incubation on ice. After washes, 4:1 ratio of Pronase E was added to the packed animals’ pellet and animals were incubated rotating at 20 °C for 5 min, followed by 12 min of gentle homogenization (2mL dounce homogenizer, pestle clearance 0.0005-0.0025 inches, DWK Life Sciences). After washes with ice-cold egg buffer (1.18 M NaCl; 480 mM KCl; 20 mM CaCl_2_; 20 mM MgCl_2_; 250 mM HEPES, pH 7.3) to remove Pronase E, cells were filtered through a 5μM filter to remove undigested animal fragments, and immediately sorted by FACS.

GLR glia cell sorting was done using a BD FACS Aria sorter equipped with 488 nm laser (Rockefeller University Flow Cytometry Resource Center), with egg buffer as the sheath buffer to preserve cell viability. Dead cell exclusion was carried out using DAPI, while DRAQ5 was used to distinguish nucleated cells from non-nucleated cell fragments. Gates for size and granularity were adjusted to exclude cell aggregates and debris. Gates for fluorescence were established using wild type (N2) nonfluorescent animals. 500,000– 800,000 YFP-positive events were sorted per replicate, which represented 0.3−0.7% of total events (after scatter exclusion), which is roughly the expected labeled-cell frequency in the animal (∼0.6%). YFP-negative events from the same gates of size and granularity, representing all other cell types, were also sorted for comparison. Cells were sorted directly into TRIzol LS (Thermo Fisher Scientific) at a ratio 3:1 (TRIzol to cell volume).

### RNAi extraction and sequencing

RNA was extracted from the sorted cells following the TRIzol LS protocol guidelines, until the isopropanol precipitation step, then RNA was re-suspended in extraction buffer of a RNA isolation kit (PicoPure, Arcturus) and isolation continued according to the manufacturer’s guideline. This two-step purification protocol helps obtaining RNA of high quality when starting with samples of large volumes, and resulted in a yield of around 10−80 ng per replica with RNA integrity number (RIN) ≥ 8, as measured by a Bioanalyzer (Agilent). All subsequent steps were performed by the Rockefeller University Genomics Resource Center. Briefly, mRNA amplification and cDNA preparation were performed using the SMARTer mRNA amplification kit (Takara #634940). Labeled samples were sequenced using an Illumina NextSeq 500 sequencer using 75 base pair single read protocols.

### RNA-seq quality assessment and differential expression analysis

Fastq files were generated with CASAVA v1.8.2 (illumina), and examined using the FASTQC (http://www.bioinformatics.babraham.ac.uk/projects/fastqc/) application for sequence quality. Reads were aligned to customized build genome that combine *C. elegans* WS262 genome release (https://downloads.wormbase.org/releases/WS262/species/c_elegans/PRJNA13758/) and the “*nep-2prom7::nls::::yfp::unc-54 3’UTR*” transgene using the STAR v2.3 aligner with parameters ^72^ (--out-FilterMultimapNmax 10 --outFilterMultimapScoreRange 1). Mapping rate was >72% with >62 million uniquely mapped reads. The alignment results were evaluated through RNA-SeQC v1.17 to make sure all samples had a consistent alignment rate and no obvious 5′ or 3′ bias ^73^. Aligned reads were summarized through featureCounts^74^ with gene models from Ensemble (Caenorhabditis_elegans. WBcel235.77.gtf) at gene level unstranded: specifically, the uniquely mapped reads (NH “tag” in bam file) that overlapped with an exon (feature) by at least 1 bp on either strand were counted and then the counts of all exons annotated to an Ensemble gene (meta features) were summed into a single number. rRNA genes, mitochondrial genes and genes with length <40 bp were excluded from downstream analysis.

Experiment was done with four independent replicates. DESeq2 was applied to normalize count matrix and to perform differential gene-expression analysis, comparing RNA counts derived from the GLR glia cells (YFP positive) to RNA counts that were derived from all other *C. elegans* cells, using negative binomial distribution ^75^.

### Microscopy

Animals were anesthetized using 100mM NaN_3_ (sodium azide) and mounted on 5% agarose pads on glass slides. Images were taken using a Zeiss compound microscope (Axio Imager M2) using MicroManager software (version 1.4.22)^76^. ImageJ ^77^ was used to produce maximum projections of z-stack images (∼0.7μm thick) presented in the Figures. Figures were prepared by using Adobe Illustrator.

### Quantification and Statistical Analysis

All microscopy fluorescence quantifications were done in ImageJ ^77^. Mutant and control animals were imaged during the same imaging session with all acquisition parameters maintained constant between the two groups. Fluorescence intensity of gene expression in GLR glia and the HMC cell (Figs. 4E, 4F, 5A, 5B, 5D, 7G) was measured in the plane with strongest signal within the z-stack in a region drawn around the GLR glia nucleus (for nuclear reporters) or cell body (for cytoplasmic reporters). A single circular region in an adjacent area was used to measure background intensity for each animal; this value was then subtracted from the fluorescence intensity of reporter expression for each GLR glia/HMC cell. Quantification of the length of the GLR glia anterior process (Fig. 7E) was performed in maximum intensity projections. A line was drawn along the anterior process for each GLR glia cell and its length was measured and normalized to the length of the pharynx for each animal. Quantification of number of puncta for *inx-18::gfp* CRISPR knock in reporter (Fig. 5C and 6E), quantification of number or percentage of GLR glia or HMC with reporter expression (Figs. 2C and D, Appendix Fig S2A and B, Fig 3A-G, Fig. 6A-E, Fig. EV2C-G, Fig. 7D, Fig. EV3A-G, and EV3J-K, Fig. 8B and D, Fig. EV4C, E and F), quantification of animals with anteriorly displaced nerve ring (Fig. 2E) and quantification of number of head muscle cells (Fig. 2F, Appendix Fig. S2C, Appendix Fig. S3A) was performed by manual counting using ImageJ.

Prism (GraphPad) was used for statistical analysis as described in Figure legends. Unpaired two-sided Students’ t test was used to determine the statistical significance between two groups.

### List of strains used and generated in this study

| Strain Name | Genotype | Comment | Reference |
| --- | --- | --- | --- |
| N2 | wild type |  | CGC |
| VC706 | *let-381(gk302) I / hT2 (I;III)* |  | 78 |
| KR429 | *dpy-5(e61) let-381(h107) unc-13(e450) I ; sDp2 (I;f)* |  | 79 |
| CB845 | *unc-30(e191) IV* |  | 63 |
| VC295 | *unc-30(ok613) IV* |  | 78 |
| GR1373 | *eri-1(mg366) IV* |  | 71 |
| CX3198 | *sax-3(ky123) X* |  | 48 |
| VC636 | *cwn-2(ok895) IV* |  | 78 |
| NG2615 | *cam-1(gm122) II* |  | 80 |
| DE60 | *dnls13 [gly-18prom::gfp + unc-119(+)] I ; unc-119(e2498) III* |  | 23 |
| MT20492 | *Lin-15B&lin-15A(n765) X ; nls471 [lgc-55prom::gfp + lin-15(+)]* |  | 21 |
| OH13025 | *otls567 [unc-46(fosmid)::SL2::H2B::mCHOPTI) + pha-1(+)]* |  | 17 |
| OH13027 | *otls569 [snf-11(fosmid)::SL2::H2B::mChopti + pha-1(+)]* |  | 17 |
| PD4443 | *ccls4443 [arg-1prom::gfp + dpy-20(+)] IV* |  | 81 |
| SD1633 | *ccls4251 [(pSAK2) myo-3p::GFP::LacZ::NLS + (pSAK4) myo-3p::mitochondrial GFP + dpy-20(+)] I ; stls10539 [dmd-4p::HIS-24::mCherry + unc-119(+)]* |  | 82 |
| OH16770 | *ccls4443 [arg-1prom::gfp + dpy-20(+)] IV ; him-5 (e1490) V ; dmd-4(ot933) X* |  | 45 |
| OP185 | *unc-119 (ed3) III ; wglS185 [fkh-2-1::TY1::EGFP::3xFLAG(92C12) + unc-119(+)]* |  | 83 |
| BC10849 | *dpy-5(e907) I ; sls10707 [sre-6prom::GFP + dpy-5(+)]* |  | 84 |
| OP447 | *unc-119(tm4063) III ; wglS447 [tag-68::TY1::EGFP::3xFLAG + unc-119(+)]* |  | 83 |
| OH9545 | *otls287 [rab-3prom1::2xnsf::yfp, rol-6(su1006)] IV* |  | 70 |
| OH441 | *otls45 [unc-119prom::gfp]* |  | 85 |
| CF4587 | *mulS253 [ eft-3prom::sfGFP1-10::unc-54 3'UTR + Cbr-unc-199(+)] II ; unc-119(ed3) III* |  | 65 |
| JN332 | *peEx3332 [nep-2prom(S)::Venus, rol6(su1006)]* |  | 20 |
| OS11153 | *nsEx5558 [nep-2prom1::GFP, pha-1(+)] line1 ; pha-1 (e2123) III* |  | This study |
| OS11157 | *nsEx5562 [nep-2prom2::GFP, pha-1(+)]line1 ; pha-1 (e2123) III* |  | This study |
| OS11158 | *nsEx5563 [nep-2prom3::GFP, pha-1(+)] line 1 ; pha-1 (e2123) III* |  | This study |
| OS11162 | *nsEx5567 [nep-2prom4::GFP, pha-1(+)] line 1 ; pha-1 (e2123) III* |  | This study |
| OS11163 | *nsEx5568 [nep-2prom5::GFP, pha-1(+)] line 1 ; pha-1 (e2123) III* |  | This study |
| OS11165 | *nsEx5570 [nep-2prom6::GFP, pha-1(+)] line 1 ; pha-1 (e2123) III* |  | This study |
| OS11166 | *nsEx5571 [nep-2prom7::GFP, pha-1(+)] line 1 ; pha-1 (e2123) III ; otls356 [rab-3prom1::NLS-TagRFP] V* |  | This study |
| OS11176 | *nsEx5581 [lgc-55prom1::GFP, pha-1(+)] line 1 ; pha-1 (e2123) III* |  | This study |
| OS11177 | *nsEx5582 [lgc-55prom2::GFP, pha-1(+)] line 1 ; pha-1 (e2123) III* |  | This study |
| OS11178 | *nsEx5583 [lgc-55prom3::GFP, pha-1(+)] line 1 ; pha-1 (e2123) III* |  | This study |
| OS11180 | *nsEx5585 [lgc-55prom4::GFP, pha-1(+)] line 1 ; pha-1 (e2123) III* |  | This study |
| OS11181 | *nsEx5586 [lgc-55prom5::GFP, pha-1(+)] line 1 ; pha-1 (e2123) III* |  | This study |
| OS11182 | *nsEx5587 [lgc-55prom6::GFP, pha-1(+)] line 1 ; pha-1 (e2123) III* |  | This study |
| OS11183 | *nsEx5588 [lgc-55prom7::GFP, pha-1(+)] line 1 ; pha-1 (e2123) III* |  | This study |
| OS11017 | *nsEx5824 [egl-6prom1::GFP, pha-1(+)] line 1 ; pha-1 (e2123) III* |  | This study |
| OS11232 | *nsEx5825 [egl-6prom2::GFP, pha-1(+)] line 1 ; pha-1 (e2123) III* |  | This study |
| OS11234 | *nsEx5827 [egl-6prom3::GFP, pha-1(+)]*<br>*line 1 ; pha-1 (e2123) III* |  | This study |
| OS11236 | *nsEx5829 [egl-6prom4::GFP, pha-1(+)]*<br>*line 1 ; pha-1 (e2123) III* |  | This study |
| OS11202 | *nsEx5591 [gly-18prom1::GFP, pha-1(+)]*<br>*line 1 ; pha-1 (e2123) III* |  | This study |
| OS11203 | *nsEx5592[gly-18prom2::GFP, pha-1(+)]*<br>*line 1 ; pha-1 (e2123) III* |  | This study |
| OS11204 | *nsEx5593 [gly-18prom3::GFP, pha-1(+)]*<br>*line 1 ; pha-1 (e2123) III* |  | This study |
| OS11205 | *nsEx5594 [gly-18prom4::GFP, pha-1(+)]*<br>*line 1 ; pha-1 (e2123) III* |  | This study |
| OS11206 | *nsEx5595 [gly-18prom5::GFP, pha-1(+)]*<br>*line 1 ; pha-1 (e2123) III* |  | This study |
| OS11208 | *nsEx5597 [gly-18prom6::GFP, pha-1(+)]*<br>*line 1 ; pha-1 (e2123) III* |  | This study |
| OS11209 | *nsEx5598 [gly-18prom7::GFP, pha-1(+)]*<br>*line 1 ; pha-1 (e2123) III* |  | This study |
| OS11211 | *nsEx5809 [gly-18prom8::GFP, pha-1(+)]*<br>*line 1 ; pha-1 (e2123) III* |  | This study |
| OS11212 | *nsEx5810 [gly-18prom9::GFP, pha-1(+)]*<br>*line 1 ; pha-1 (e2123) III ; otIs356 [rab-*<br>*3prom1::NLS-TagRFP] V* |  | This study |
| OS11168 | *nsEx5573 [hlh-1prom1::GFP, pha-1(+)]*<br>*line 1 ; pha-1 (e2123) III* |  | This study |
| OS11171 | *nsEx5576 [hlh-1prom2::GFP, pha-1(+)]*<br>*line 1 ; pha-1 (e2123) III* |  | This study |
| OS11172 | *nsEx5577 [hlh-1prom3 cloned in pSM-*<br>*GFP, pha-1(+)] line 1 ; pha-1 (e2123) III* |  | This study |
| OS11959 | *nsEx6055 [inx-18prom1::GFP, pha-1(+)]*<br>*line 1 ; pha-1 (e2123) III* |  | This study |
| OS11960 | *nsEx6056 [inx-18prom2::GFP, pha-1(+)]*<br>*line 1 ; pha-1 (e2123)* |  | This study |
| OS11962 | *nsEx6058 [inx-18prom3::GFP, pha-1(+)]*<br>*line 1 ; pha-1 (e2123) III* |  | This study |
| OS12795 | *nxEx6276 [inx-18prom4::gfp + pha-1+]*<br>*line 1 ; pha-1 (e2123) III* |  | This study |
| OS12797 | *nxEx6278 [inx-18prom5::gfp + pha-1+]*<br>*line 1 ; pha-1 (e2123) III* |  | This study |
| OS12798 | *nxEx6279 [inx-18prom6::gfp + pha-1+]*<br>*line 1 ; pha-1 (e2123) III* |  | This study |
| OS12800 | *nxEx6281 [inx-18prom7::gfp + pha-1+]*<br>*line 1 ; pha-1 (e2123) III* |  | This study |
| OS12801 | *nxEx6282 [inx-18prom8::gfp + pha-1+]*<br>*line 1 ; pha-1 (e2123) III* |  | This study |
| OS12802 | *nxEx6283 [inx-18prom9::gfp + pha-1+]*<br>*line 1 ; pha-1 (e2123) III* |  | This study |
| OS12819 | *nsEx6288 [inx-18prom10::gfp, pha-1+]*<br>*line 1 ; pha-1(e2123) III* |  | This study |
| OS12821 | *nsEx6290 [inx-18prom11::gfp, pha-1+]*<br>*line 1 ; pha-1(e2123) III* |  | This study |
| OS14565 | *nsEx7189 [unc-30prom1::gfp + pha-1+]*<br>*line 1 ; pha-1(e2123) III* |  | This study |
| OS14566 | *nsEx7190 [unc-30prom2::gfp + pha-1+]*<br>*line 1 ; pha-1(e2123) III* |  | This study |
| OS14567 | *nsEx7191 [unc-30prom3::gfp + pha-1+] line 1 ; pha-1(e2123) III* |  | This study |
| OS14568 | *nsEx7192 [unc-30prom4::gfp + pha-1+] line 1 ; pha-1(e2123) III* |  | This study |
| OS14569 | *nsEx7193 [unc-30prom5::gfp + pha-1+] line 1 ; pha-1(e2123) III* |  | This study |
| OS14570 | *nsEx7194 [unc-30prom6::gfp + pha-1+] line 1 ; pha-1(e2123) III* |  | This study |
| OS14571 | *nsEx7195 [unc-30prom7::gfp + pha-1+] line 1 ; pha-1(e2123) III* |  | This study |
| OS13296 | *nsEx6463 [unc-30prom8::gfp + pha-1+] line 1 ; pha-1(e2123) III* |  | This study |
| OS14572 | *nsEx7196 [unc-30prom9::gfp + pha-1+] line 1 ; pha-1(e2123) III* |  | This study |
| OS14573 | *nsEx7197 [unc-30prom10::gfp + pha-1+] line 1 ; pha-1(e2123) III* |  | This study |
| OS14094 | *nsEx6871 [F41G4.8prom1::gfp + pha-1+] line 1 ; pha-1 (e2123) III* |  | This study |
| OS14173 | *nsEx6921 [F41G4.8prom2::gfp + pha1+] line 1 ; pha-1 (e2123) III; nsls758 [nep-2 prom7::tagrf] V* |  | This study |
| OS14164 | *nsEx6914 [twk-4prom1::gfp + pha-1+] line 1 ; pha-1 (e2123) III* |  | This study |
| OS14158 | *nsEx6908 [twk-4prom2::gfp + pha-1+] line 1 ; pha-1 (e2123) III* |  | This study |
| OS14098 | *nsEx6875 [twk-4prom3::gfp + pha-1+] line 1 ; pha-1 (e2123) III* |  | This study |
| OS14157 | *nsEx6907 [ocr-1prom1::gfp + pha-1+] line 1 ; pha-1 (e2123) III* |  | This study |
| OS14273 | *nsEx6966 [ocr-1prom2::gfp + pha-1+] line 1 ; pha-1 (e2123) III* |  | This study |
| OS14097 | *nsEx6874 [ocr-1prom3::gfp + pha-1+] line 1 ; pha-1 (e2123) III* |  | This study |
| OS14099 | *nsEx6876 [twk-9prom1::gfp +pha-1+] line 1 ; pha-1 (e2123) III* |  | This study |
| OS14172 | *nsEx6920 [mig-6prom1::gfp + pha-1+] line 1 ; pha-1 (e2123) III* |  | This study |
| OS14156 | *nsEx6906 [mig-6prom2::gfp + pha-1+] line 1 ; pha-1 (e2123) III* |  | This study |
| OS14174 | *nsEx6922 [mig-6prom3::gfp + pha-1+] line 1 ; pha-1 (e2123) III* |  | This study |
| OS14161 | *nsEx6911 [mig-6prom4::gfp + pha-1+] line 1 ; pha-1 (e2123) III* |  | This study |
| OS14175 | *nsEx6923 [mig-6prom5::gfp + pha-1+] line 1 ; pha-1 (e2123) III* |  | This study |
| OS14198 | *nsEx6941 [mig-17prom1::gfp + pha-1+] line 1 ; pha-1 (e2123) III* |  | This study |
| OS14188 | *nsEx6931 [mig-17prom2::gfp + pha-1+] line 1 ; pha-1 (e2123) III* |  | This study |
| OS14176 | *nsEx6924 [let-2prom1::gfp + pha-1+] line 1 ; pha-1 (e2123) III* |  | This study |
| OS14159 | *nsEx6909 [let-2prom2::gfp + pha-1+] line 1 ; pha-1 (e2123) III* |  | This study |
| OS14160 | *nsEx6910 [let-2prom3::gfp + pha-1+] line 1 ; pha-1 (e2123) III* |  | This study |
| OS14187 | *nsEx6930 [let-2prom4::gfp + pha-1+] line 1; pha-1 (e2123) III* |  | This study |
| OS14197 | *nsEx6940 [let-2prom5::gfp + pha-1+] line 1; pha-1 (e2123) III* |  | This study |
| OS14276 | *nsEx6969 [acc-2prom1::gfp + pha-1+] line 1; pha-1 (e2123) III* |  | This study |
| OS14268 | *nsEx6961 [oac-7prom1::gfp + pha-1+] line 1; pha-1 (e2123) III; nsls758 [nep-2 prom7::nls::yfp] V* |  | This study |
| OS14307 | *nsEx6973 [kvs-5prom1::rfp + pha-1+] line 1; pha-1 (e2123) III; nsls758 [nep-2 prom7::nls::yfp] V* |  | This study |
| OS14203 | *nsEx6942 [kvs-5prom2::rfp + pha-1+] line 1; pha-1 (e2123) III; nsls758 [nep-2 prom7::nls::yfp] V* |  | This study |
| OS14272 | *nsEx6965 [lbp-1prom1::rfp + pha-1+] line 1; pha-1 (e2123) III; nsls758 [nep-2 prom7::nls::yfp] V* |  | This study |
| OS14279 | *nsEx6972 [lbp-1 prom2::rfp + pha-1+] line 1; pha-1 (e2123) III; nsls758 [nep-2 prom7::nls::yfp] V* |  | This study |
| OS14308 | *nsEx6974 [adt-3prom1::rfp + pha-1+] line 1; pha-1 (e2123) III; nsls758 [nep-2 prom7::nls::yfp] V* |  | This study |
| OS14343 | *nsEx7015 [R03E9.2 prom1::rfp + pha-1+] line 1; pha-1 (e2123) III; nsls758 [nep-2 prom7::nls::yfp] V* |  | This study |
| OS14455 | *nsEx7118 [R03E9.2prom 2::rfp + pha-1+] line 1; pha-1 (e2123) III; nsls758 [nep-2 prom7::nls::yfp] V* |  | This study |
| OS14345 | *nsEx7017 [R03E9.2prom3::rfp + pha-1+] line 1; pha-1 (e2123) III; nsls758 [nep-2 prom7::nls::yfp] V* |  | This study |
| OS14346 | *nsEx7018 [R03E9.2prom4::rfp + pha-1+] line 1; pha-1 (e2123) III; nsls758 [nep-2 prom7::nls::yfp] V* |  | This study |
| OS14398 | *nsEx7067 [F49B2.6prom1::rfp + pha-1+] line 1; pha-1 (e2123) III; nsls758 [nep-2 prom7::nls::yfp] V* |  | This study |
| OS14399 | *nsEx7068 [F49B2.6prom2::rfp + pha-1+] line 1; pha-1 (e2123) III; nsls758 [nep-2 prom7::nls::yfp] V* |  | This study |
| OS14395 | *nsEx7064 [nta-1prom1::rfp + pha-1+] line 1; pha-1 (e2123) III; nsls758 [nep-2 prom7::nls::yfp] V* |  | This study |
| OS14575 | *nsEx7199 [nta-1prom2::rfp + pha-1+] line 1 ; pha-1 (e2123) III ; nsls758 [nep-2 prom7::nls::yfp] V* |  | This study |
| OS14400 | *nsEx7069 [T14B4.9prom1::rfp + pha-1+] line 1; pha-1 (e2123) III; nsls758 [nep-2 prom7::nls::yfp] V* |  | This study |
| OS14458 | *nsEx7121 [T14B4.9prom2::rfp + pha-1+] line 1; pha-1 (e2123) III; nsls758 [nep-2 prom7::nls::yfp] V* |  | This study |
| OS14483 | *nsEx7138 [T14B4.9prom3::rfp + pha-1+] line 1; pha-1 (e2123) III; nsls758 [nep-2 prom7::nls::yfp] V* |  | This study |
| OS14459 | *nsEx7122 [tbc-12prom1::rfp + pha-1+] line 1; pha-1 (e2123) III; nsls758 [nep-2 prom7::nls::yfp] V* |  | This study |
| OS14574 | *nsEx7198 [tbc-12prom2::rfp + pha-1+] line 1; pha-1 (e2123) III; nsls758 [nep-2 prom7::nls::yfp] V* |  | This study |
| OS14482 | *nsEx7137 [tbc-12prom3::rfp + pha-1+] line 1; pha-1 (e2123) III; nsls758 [nep-2 prom7::nls::yfp] V* |  | This study |
| OS14484 | *nsEx7139 [tbc-12prom4::rfp + pha-1+] line 1; pha-1 (e2123) III; nsls758 [nep-2 prom7::nls::yfp] V* |  | This study |
| OS14454 | *nsEx7117 [pgp-4prom 1::rfp + pha-1+] line 1; pha-1 (e2123) III; nsls758 [nep-2 prom7::nls::yfp] V* |  | This study |
| OS14457 | *nsEx7120 [pgp-4prom 2::rfp + pha-1+] line 1; pha-1 (e2123) III; nsls758 [nep-2 prom7::nls::yfp] V* |  | This study |
| OS14205 | *nsEx6944 [F19B10.3prom1::rfp + pha-1+] line 1; pha-1 (e2123) III; nsls758 [nep-2 prom7::nls::yfp] V* |  | This study |
| OS14576 | *nsEx7200 [haf-7prom1::rfp + pha-1+] line 1; pha-1 (e2123); nsls758 [nep-2 prom7::nls::yfp] V* |  | This study |
| OS14456 | *nsEx7119 [haf-7prom 2::rfp + pha-1+] line 1; pha-1 (e2123) III; nsls758 [nep-2 prom7::nls::yfp] V* |  | This study |
| OS13444 | *nsEx6539 [glb-26prom1::gfp + pha-1+] line 1; pha-1(e2123) III* |  | This study |
| OS13445 | *nsEx6540 [glb-26prom1::gfp + pha-1+] line 2; pha-1(e2123) III* |  | This study |
| OS12140 | *nsEx6132 [WRM0641dF02 gly-18::sl2::snl::yfp::h2b, pha-1+] line1; pha-1 (e2123) III* |  | This study |
| OS12142 | *nsEx6134 [WRM065dE01 gpx-8::sl2::snl::yfp::h2b, pha-1+] line1; pha-1 (e2123) III* |  | This study |
| OS12145 | *nsEx6137[F28H7.2prom1::gfp, pha-1+] line1; pha-1 (e2123) III* |  | This study |
| OS12147 | *nsEx6139 [T28A11.3prom1::gfp, pha-1+] line 1; pha-1 (e2123) III* |  | This study |
| OS12149 | *nsEx6141[ZC317.2prom1::gfp, pha-1+] line 1; pha-1 (e2123) III* |  | This study |
| OS12151 | *nsEx6143 [F35D2.3prom1::gfp, pha-1+] line 1; pha-1 (e2123) III* |  | This study |
| OS12789 | *nsEx6078 [F13D12.10prom1::GFP, pha-1(+)] line1; pha-1 (e2123) III; nsls700 V* |  | This study |
| OS11967 | *nsEx6063 [mnp-1prom3::GFP, pha-1(+)] line 1; pha-1 (e2123) III* |  | This study |
| OS11219 | *nsEx5817 [ncam-1prom2::GFP, pha-1(+)] line1; pha-1 (e2123) III* |  | This study |
| OS11956 | *nsEx6052 [unc-54prom3::GFP, pha-1(+)] line1 ; pha-1 (e2123)* |  | This study |
| OS12370 | *nsEx6202 [WRM061BG12 best-22 TY1::EGFP::3xFLAG(92C12) + pha-1(+)] line 1; pha-1 (e2123) III* |  | Tagged fosmid generated by <sup>83</sup>. Transgenic line generated in this study. |
| OS14023 | *nsEx6821 [WRM069bF08 (let-381 fosmid) + myo-3p::mCherry] line 1 ; let-381(gk302) I ; nsls746 [nep-2prom7::gfp] V* |  | This study |
| OS14024 | *nsEx6822 [WRM069bF08 (let-381 fosmid) + myo-3p::mCherry] line 2 ; let-381(gk302) I ; nsls746 [nep-2prom7::gfp] V* |  | This study |
| OS12240 | *nsEx6151 [WRM0639aA08 (unc-30 fosmid) + unc-122prom::gfp] line 1 ; unc-30 (e191) IV ; nsls700 [nep-2prom7::tagrfp] V* |  | This study |
| OS12241 | *nsEx6152 [WRM0639aA08 (unc-30 fosmid) + unc-122prom::gfp] line 2 ; unc-30 (e191) IV ; nsls700 [nep-2prom7::tagrfp] V* |  | This study |
| OS12242 | *nsEx6153 [WRM0639aA08 (unc-30 fosmid) + unc-122prom::gfp] line 3 ; unc-30 (e191) IV ; nsls700 [nep-2prom7::tagrfp] V;* |  | This study |
| OS12246 | *nsEx6157 [WRM0617aE02 (unc-30 fosmid) + unc-122prom::rfp] line 1 ; unc-30 (e191) IV ; nsls746 [nep-2prom7::gfp] V* |  | This study |
| OS12247 | *nsEx6158 [WRM0617aE02 (unc-30 fosmid) + unc-122prom::rfp] line 2 ; unc-30 (e191) IV ; nsls746 [nep-2prom7::gfp] V* |  | This study |
| OS12244 | *nsEx6155 [unc-30 whole locus PCR + unc-122prom::gfp] line 1 ; unc-30 (e191) IV ; nsls700 [nep-2prom7::tagrfp] V* |  | This study |
| OS13665 | *nsEx6668 [nep-2p7::let-381 sas RNAi + myo-3::mCherry] line 1 ; let-381(ns995) I; eri-1(mg366) IV* |  | This study |
| OS13666 | *nsEx6669 [nep-2p7::let-381 sas RNAi + myo-3::mCherry] line 2; let-381(ns995) I; eri-1(mg366) IV* |  | This study |
| OS13667 | *nsEx6670 [nep-2p7::let-381 sas RNAi + myo-3::mCherry] line 3; let-381(ns995) I; eri-1(mg366) IV* |  | This study |
| OS13668 | *nsEx6671 [nep-2p7 (sas RNAi control) + myo-3::mCherry] line 1; let-381(ns995) I; eri-1(mg366) IV* |  | This study |
| OS13820 | *nsEx6667 [nep-2p7 (sas RNAi control) + myo-3::mCherry] line 2 ; nep-2(syb4689) II; eri-1(mg366) IV* |  | This study |
| OS13672 | *nsEx6675 [nep-2p7 (sas RNAi control) + myo-3::mCherry] line 3; ccls4443 [arg-1prom::gfp + dpy-20(+)] eri-1(mg366) IV* |  | This study |
| OS13435 | *nsEx6530 [hsp-16,2prom::let-381 cDNA, hsp-16,2prom::unc-30 cDNA, pha-1+] line 1 ; pha-1(e2123) III ; nsls700 [nep-2prom7::tagrfp] V* |  | This study |
| OS13436 | *nsEx6531 [hsp-16,2prom::let-381 cDNA, hsp-16,2prom::unc-30 cDNA, pha-1+] line 2 ; pha-1(e2123) III ; nsls700 [nep-2prom7::tagrfp] V* |  | This study |
| OS13438 | *nsEx6533 [hsp-16,2prom::let-381 cDNA, pha-1+] line 1 ; pha-1(e2123) III ; nsls700 [nep-2prom7::tagrfp] V* |  | This study |
| OS13439 | *nsEx6534 [hsp-16,2prom::let-381 cDNA, pha-1+] line 2 ; pha-1(e2123) III ; nsls700 [nep-2prom7::tagrfp] V* |  | This study |
| OS13440 | *nsEx6535 [hsp-16,2prom::unc-30 cDNA, pha-1+] line 1 ; pha-1(e2123) III ; nsls700 [nep-2prom7::tagrfp] V* |  | This study |
| OS13441 | *nsEx6536 [hsp-16,2prom::unc-30 cDNA, pha-1+] line 2 ; pha-1(e2123) III ; nsls700 [nep-2prom7::tagrfp] V* |  | This study |
| OS11484 | *nsls700 [nep-2prom7::tagrfp] V* |  | This study |
| OS11703 | *nsls746 [nep-2prom7::gfp] V* |  | This study |
| OS11715 | *nsls758 [nep-2prom7::nls::yfp] V* |  | This study |
| OS12099 | *nsls831 [pll-1prom1::tagrfp] X* |  | This study |
| OS12103 | *nsls835 [hot-5prom2::gfp] X* |  | This study |
| OS12767 | *nsls879 [nep-2p7::tir-1::mCardinal + unc-122prom::mCherry]* |  | This study |
| OS12156 | *nsls854 [nep-2prom7::egl-1, myo-3mCherry] ; nsls831 [pll-1prom1::tagrfp] X* |  | This study |
| OS12700 | *unc-30(ns959[unc-30::gfp::degron(AID)]) IV* |  | This study |
| OS13288 | *let-381(ns995[let-381::gfp::aid]) I* |  | This study |
| PHX4689 | *nep-2(syb4689[gfp::h2b::sl2::nep-2]) II* |  | This study |
| PHX3025 | *hlh-1(syb3025[hlh-1::gfp::aid]) II* |  | This study |
| PHX2879 | *inx-18(syb2879[inx-18::aid::emgfp]) IV* |  | This study |
| PHX5792 | *pll-1(syb5792[pll-1::sl2::gfp::h2b]) III* |  | This study |
| PHX5704 | *gbb-1(syb5704[gbb-1::sl2::gfp::h2b]) X* |  | This study |
| PHX5759 | *gbb-2(syb5759[gbb-2::sl2::gfp::h2b]) IV* |  | This study |
| OS13744 | *nep-2(ns1012[\*syb4689]) II* | *ns1012 = let-381 motif mutation* | This study |
| OS14214 | *pll-1(ns1040[\*syb5792]) III* | *ns1040 = let-381 motif mutation* | This study |
| OS13748 | *hlh-1(ns1016[\*syb3025]) II* | *ns1016 = 1<sup>st</sup> let-381 motif mutation* | This study |
| OS13842 | *hlh-1(ns1027[\*syb3025]) II* | *ns1027 = 2<sup>nd</sup> let-381 motif mutation* | This study |
| OS13747 | *hlh-1(ns1015[\*syb3025]) II* | *ns1015 = 1<sup>st</sup> + 2<sup>nd</sup> motif mutation* | This study |
| OS13742 | *inx-18(ns1010[\*syb2879])* | *ns1010 = let-381 motif mutation* | This study |
| OS13835 | *let-381(ns1023[\*ns995]) I* | *ns1023 = let-381 motif deletion type 1* | This study |
| OS13838 | *let-381(ns1026[\*ns995]) I* | *ns1026 = let-381 motif deletion type 2* | This study |
| OS13308 | *unc-30(ns1000[\*ns959]) IV* | *ns1000 = 1<sup>st</sup> let-381 motif mutation* | This study |
| OS13307 | *unc-30(ns999[\*ns959]) IV* | *ns999 = 1<sup>st</sup> + 2<sup>nd</sup> let-381 motifs mutation* | This study |
| OS13309 | *unc-30(ns1001[\*ns959]) IV* | *ns1001 = 3<sup>rd</sup> let-381 motif mutation* | This study |
| OS13306 | *unc-30(ns998[\*ns959])* | *ns998 = three let-381 motifs deletion* | This study |

## AUTHOR CONTRIBUTIONS

**Nikolaos Stefanakis**.: Conceptualization; Methodology; Formal Analysis; Investigation; Writing – Original Draft; Writing – Review and Editing; Visualization; Funding Acquisition. **Jessica Jiang**: Investigation. **Yupu Liang**: Software; Formal Analysis. **Shai Shaham**: Conceptualization; Writing – Original draft; Writing – Review and Editing; Project Administration; Funding Acquisition.

## ACKNOWLEDGEMENTS

We thank Oliver Hobert and Yuichi Lino for strains; The Rockefeller University Flow Cytometry and Genomics Resource Centers for technical support; Menachem Katz and members of the Shaham lab for experimental advice, comments and discussion. Some strains were provided by the CGC, which is funded by NIH Office of Research Infrastructure Programs (P40 OD010440). This work was supported in part by funds from a Leon Levy Fellowship to N.S. and by NIH grant R35NS105094 to S.S.

---

**Figure EV1.**
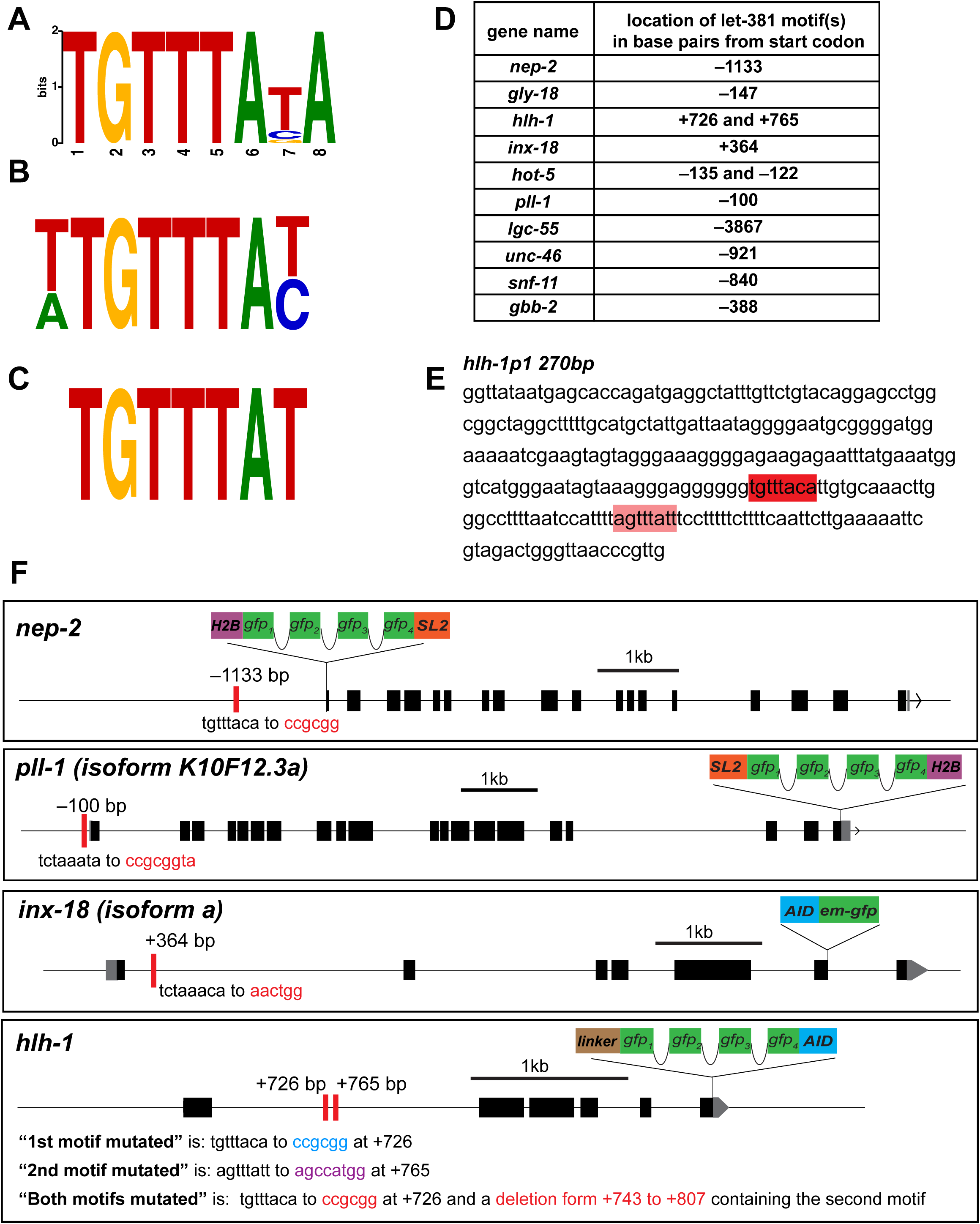
*let-381* motifs are required for endogenous GLR gene expression and *let-381* autoregulation in GLR glia. (A) *let-381* motif identified in this study. (B) Motif of the *let-381* ortholog *foxf* from ^40^. (C) *let-381* motif from ^41^. Similarities between the three motifs are apparent. (D) Locations (distances from start codons) of *let-381* motifs of genes whose expression in GLR is downregulated in *let-381* mutants (either GLR-specific *let-381* RNAi and/or the *let-381* autoregulatory allele). (E) Minimal promoter *hlh-1prom1* was one of the promoters used in MEME to identify common motifs present in GLR glia genes. The *let-381* motif identified by MEME is highlighted in dark red. A *let-381* motif with slightly altered sequence (light red) was identified manually later and is required, together with the first motif, to control *hlh-1* expression in GLR glia. (F) Schematics showing details on endogenous *gfp*-based tags, location of *let-381* motifs and their mutation for *nep-2*, *pll-1*, *hlh-1* and *inx-18* genes. Red bars represent *let-381* motifs. Distance from ATG is indicated above each motif. Nucleotide changes for each motif mutation is shown below the motifs.

**Figure EV2.**
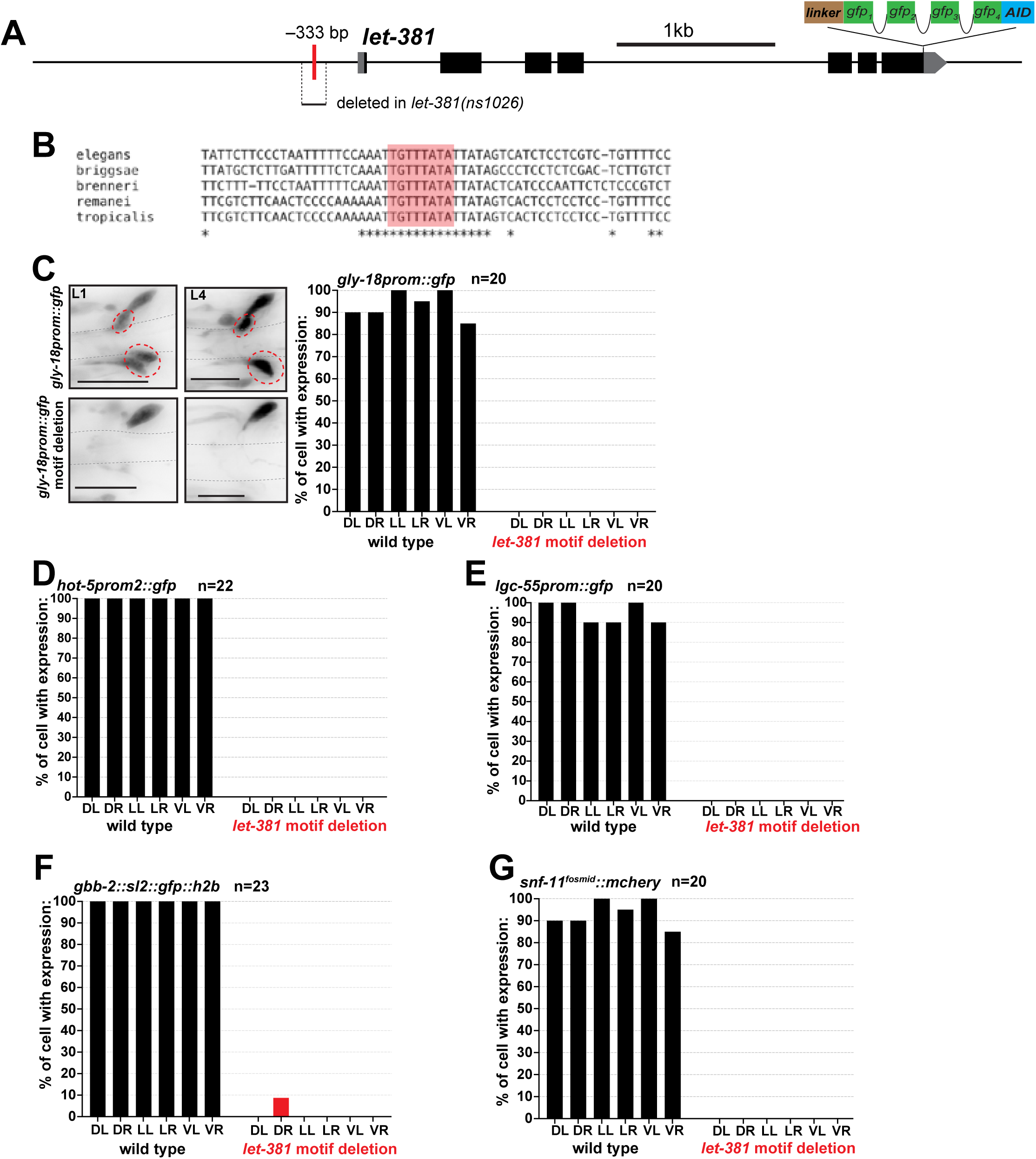
GLR gene expression is lost in *let-381* autoregulatory mutant animals. (A) Schematic showing the location of the *let-381* motif (red bar) in the *let-381* promoter region and region deleted in the *let-381(ns1026)* mutation. (B) Conservation of the *let-381* autoregulatory motif sequence (red box) is shown among five nematode species. Asterisks indicate conserved nucleotides. (C-G) Effect of *let-381* autoregulatory motif deletion on expression of (C) *gly-18*, (D) *hot-5*, (E) *lgc-55*, (F) *gbb-2* and (G) *snf-11* in GLR glia. Bar graphs show quantifications of gene expression at the L4 stage. For (C) animal images showing gene expression at L1 and L4 stages in wildtype and mutant backgrounds are shown on the left. Dashed red circles outline expression in GLR glia. Data information: Anterior is left, dorsal is up and scale bars are 10 μm for all animal images.

**Figure EV3.**
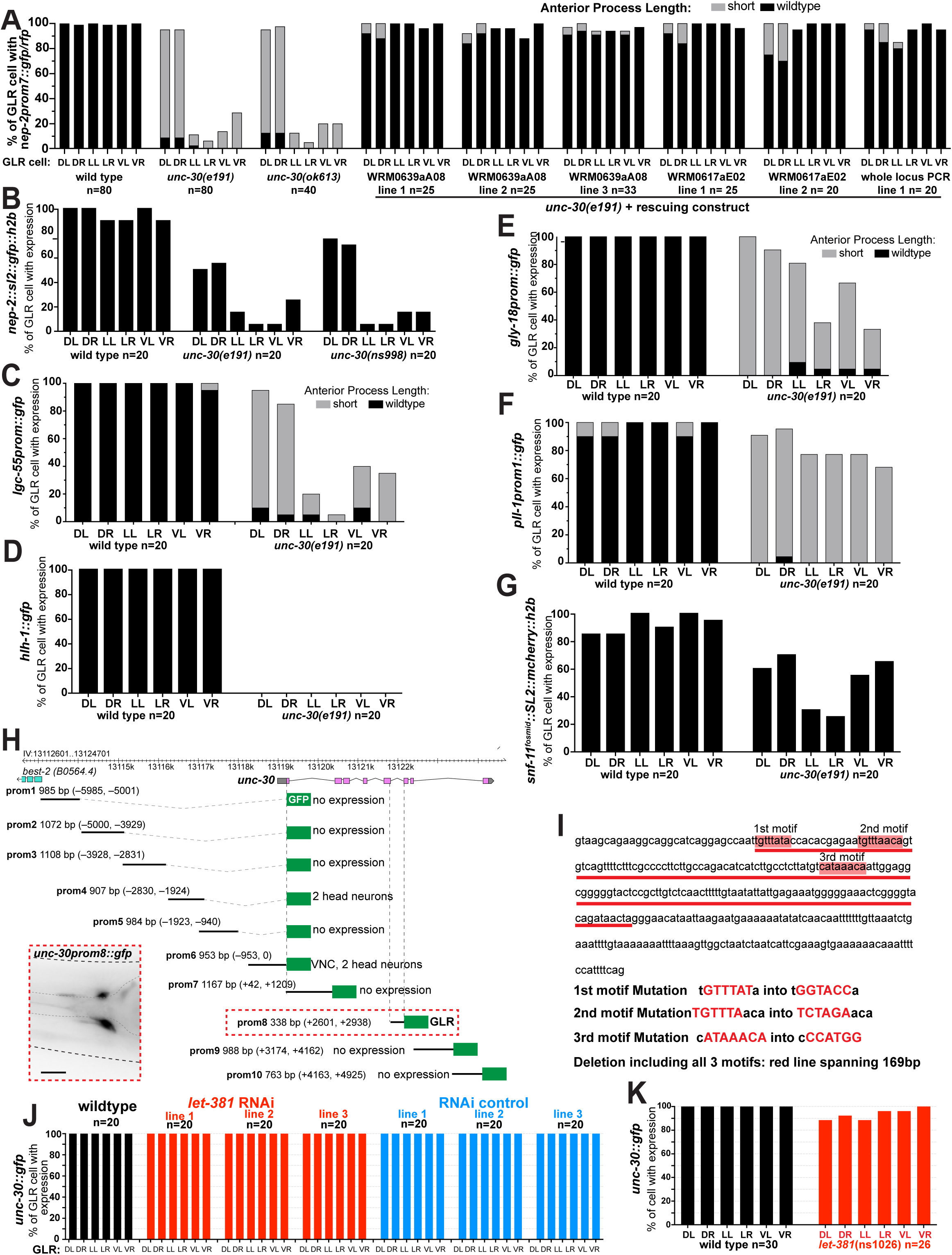
Effect of *unc-30* on GLR gene expression. (A) Transgenic constructs containing different fosmid clones (WRM) or PCR amplicons carrying wild type copies of UNC-30 can rescue the effect of *unc-30(e191)* on GLR gene expression and anterior process length. (B-G) Effect of *unc-30* mutation on expression of different genes in GLR glia. Expression of (E) *gly-18*, (F) *pll-1* and (G) *snf-11* is affected at a lesser extent compared to (B) *nep-2*, (C) *lgc-55* and (D) *hlh-1*. (H) Cis-regulatory dissection analysis of *unc-30*. The fifth intron (prom8) of *unc-30* is sufficient to drive expression in GLR glia. (I) Three *let-381* motifs are found in the fifth intron of *unc-30* (red boxes). Details on *let-381* motif mutation alleles are shown below the DNA sequence. (J), (K) Endogenous *unc-30::gfp* expression is not affected by postembryonic *let-381* knockdown either (J) by GLR-specific RNAi or (K) in the GLR-specific *let-381* autoregulatory motif deletion allele *let-381(ns1026)*. Data information: Anterior is left, dorsal is up and scale bars are 10 μm for all animal images.

**Figure EV4.**
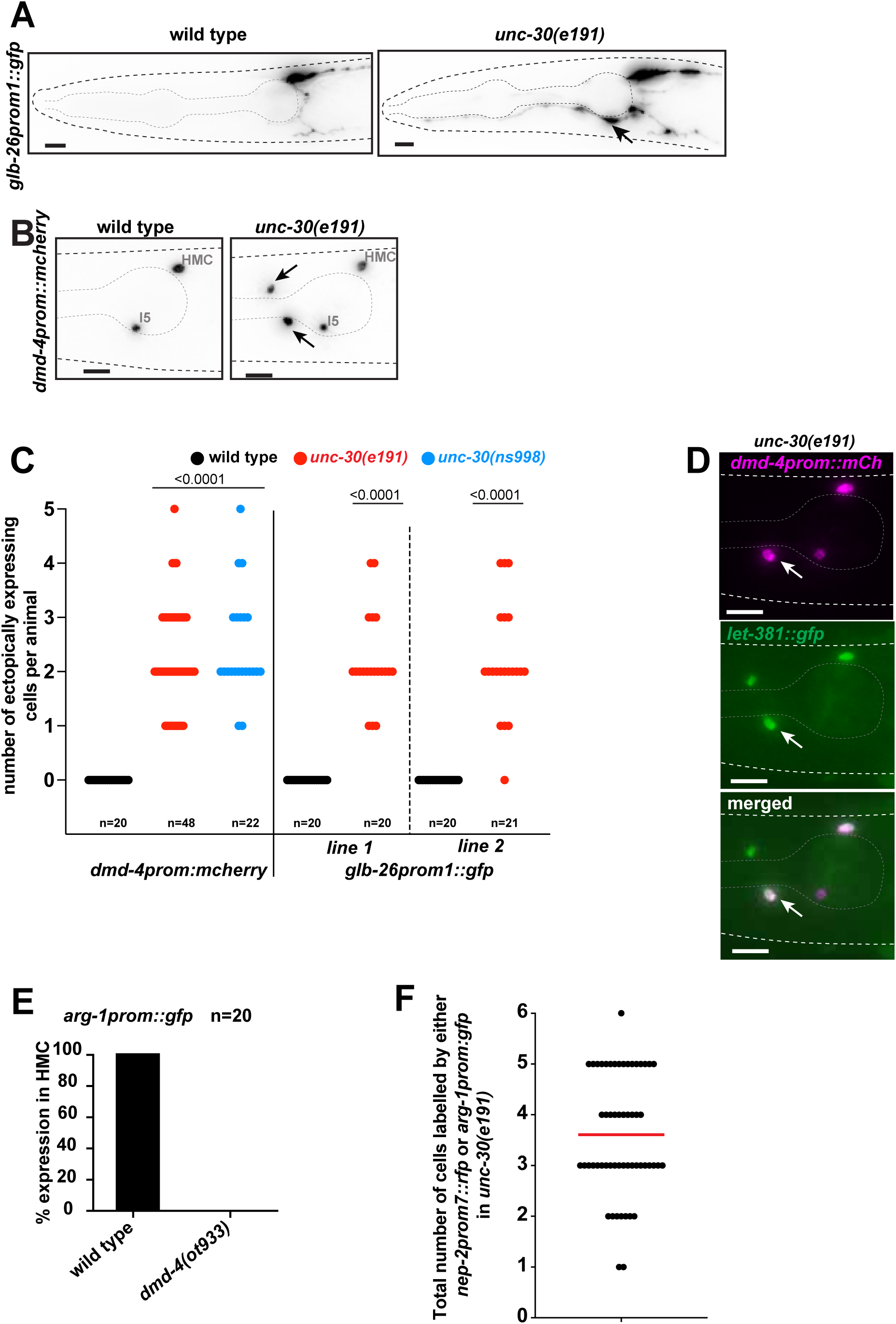
*unc-30* represses HMC gene expression in GLR glia. (A) Fluorescence images showing expression of *glb-26prom::gfp* in wild type and *unc-30(e191)* mutants. (B) Fluorescence images showing expression of *dmd-4prom::mCherry* in wild type and *unc-30(e191)* mutants. Arrows point to ectopic expression in *unc-30(e191)* mutants. (C) Quantification of ectopic expression of the two HMC reporters shown in (A) and (B) in *unc-30* mutant backgrounds. (D) Cells ectopically expressing (white arrow) the HMC reporter *dmd-4prom::mCherry* (magenta) always co-express *let-381::gfp* (green). (E) Expression of *arg-1prom::gfp* is lost in HMC in *dmd-4(ot933)* mutants. (F) Total number of GLR glia cells expressing either the GLR glia-specific *nep-2prom7::rfp* or the HMC-specific *arg-1prom::gfp* in *unc-30(e191)* mutants. Data information: Un-paired t-test used for statistical analysis in (C). Anterior is left, dorsal is up and scale bars are 10 μm for all animal images.

**Figure EV5.**
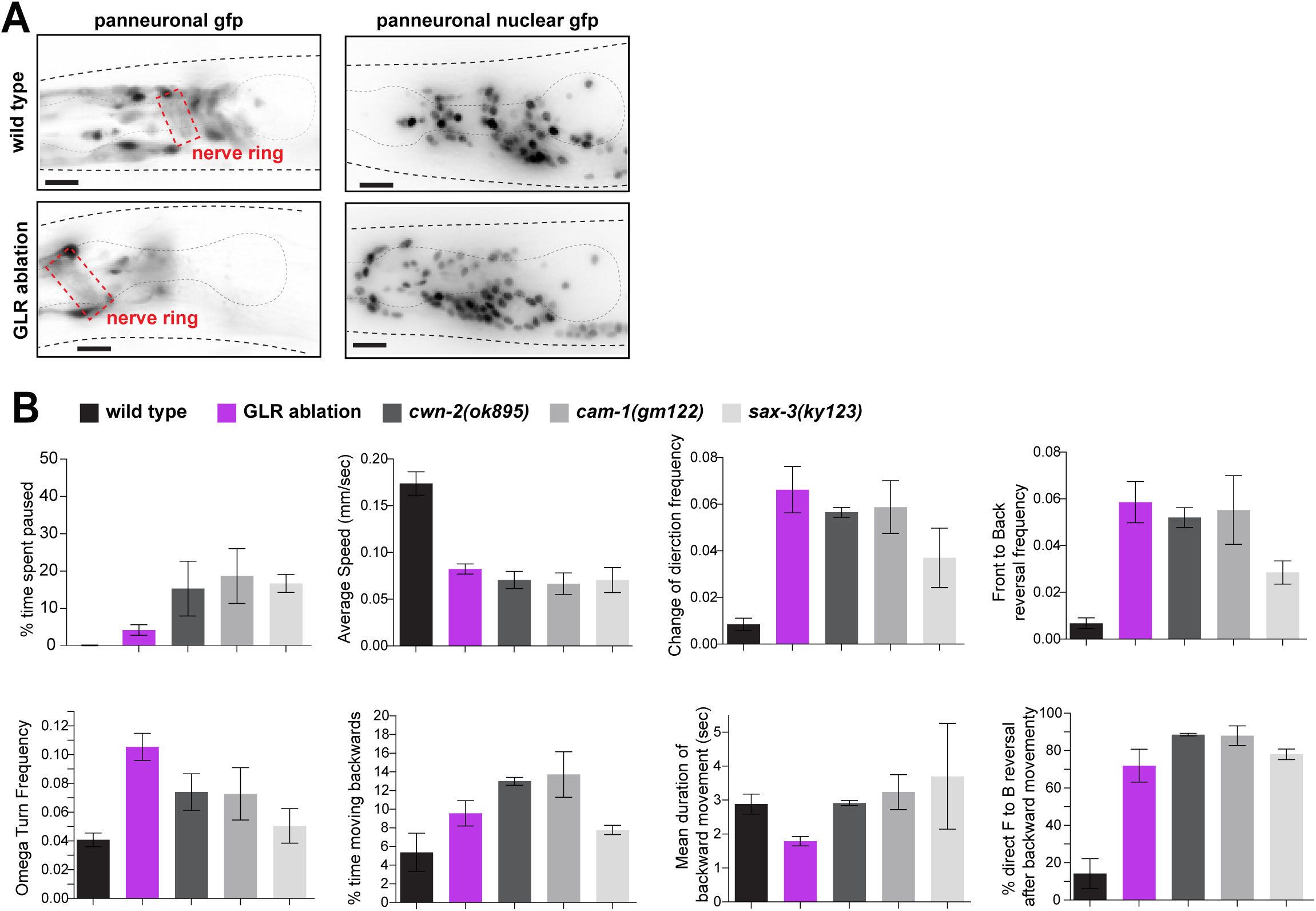
Locomotion defects of GLR-ablated animals could partially be due to anteriorly displaced nerve ring. (A) In wild type animals (top row), axons of the nerve ring (dashed red box) are located between the two pharyngeal bulbs. In GLR ablated animals (bottom row) the nerve ring is anteriorly displaced, located on top of the anterior pharynx bulb. As evidenced in the images on the right, not only the axonal projections, but also neuronal cell bodies (panneuronal nuclear gfp) are anteriorly displaced. Panneuronal gfp = *unc-119prom::gfp*, panneuronal nuclear gfp = *rab-3prom1::nls::yfp*. (B) *cwn-2(ok895)*, *cam-1(gm122)*, *sax-3(ky123)* mutants with anteriorly displaced nerve rings exhibit locomotion defects to the same direction, although of different magnitude as the GLR glia-ablated animals. Data information: Two to four movies were analyzed per genotype for *cwn-2*, *cam-1* and *sax-3* mutants in (B). Error bars show standard deviation in (B). Anterior is left, dorsal is up and scale bars are 10 μm for all animal images.

**Table EV1.** *let-381* motif-containing minimal promoters from 14 out of 19 genes are sufficient to drive expression of these genes in GLR glia. Promoters driving GLR glia expression are shown in green. Promoters not driving GLR glia expression are shown in yellow. Three independent extrachromosomal arrays were scored for each promoter. Computationally predicted motifs are not always functional in vivo^86^, which explains why only 20 out of 37 total promoters drive GLR glia expression.

| GLR enriched gene | gfp-based promoter fusion | promoter size (bp) | # of *let-381* motifs in promoter | Location of motifs from start codon |
| --- | --- | --- | --- | --- |
| *F41G4.8* | *promoter 1::gfp* | 220 | 2 | −315, −212 |
|  | *promoter 2::gfp* | 149 | 1 | −1661 |
| *twk-4* | *twk-4 promoter 1::gfp* | 159 | 1 | −3634 |
|  | *twk-4 promoter 2::gfp* | 172 | 1 | −1320 |
|  | *twk-4 promoter 3::gfp* | 167 | 1 | −384 |
| *ocr-1* | *ocr-1 promoter 1::gfp* | 165 | 1 | −996 |
|  | *ocr-1 promoter 2::gfp* | 162 | 1 | −242 |
|  | *ocr-1 promoter 3::gfp* | 169 | 1 | +924 |
| *twk-9* | *twk-9 promoter 1::gfp* | 214 | 1 | −62 |
| *mig-6* | *mig-6 promoter 1::gfp* | 168 | 1 | −1561 |
|  | *mig-6 promoter 2::gfp* | 170 | 1 | −1149 |
|  | *mig-6 promoter 3::gfp* | 181 | 1 | −372 |
|  | *mig-6 promoter 4::gfp* | 193 | 1 | +2128 |
|  | *mig-6 promoter 5::gfp* | 187 | 1 | +9758 |
| *mig-17* | *mig-17 promoter 1::gfp* | 177 | 1 | −193 |
|  | *mig-17 promoter 2::gfp* | 162 | 1 | +273 |
|  | *mig-17 promoter 3::gfp* | 171 | 2 | +415, + 508 |
| *let-2* | *let-2 promoter 1::gfp* | 174 | 1 | −2780 |
|  | *let-2 promoter 2::gfp* | 149 | 2 | −1965, −1944 |
|  | *let-2 promoter 3::gfp* | 214 | 1 | −500 |
|  | *let-2 promoter 4::gfp* | 177 | 1 | +611 |
|  | *let-2 promoter 5::gfp* | 184 | 1 | +1550 |
| *acc-2* | *acc-2 promoter 1::gfp* | 209 | 2 | −1446, −1389 |
| *oac-7* | *oac-7 promoter 1::rfp* | 180 | 1 | +2561 |
| *kvs-5* | *kvs-5 promoter 1::rfp* | 200 | 1 | +867 |
|  | *kvs-5 promoter 2::rfp* | 152 | 1 | +3138 |
| *lbp-1* | *lbp-1 promoter 1::rfp* | 158 | 1 | −1090 |
|  | *lbp-1 promoter 2::rfp* | 163 | 1 | +491 |
| *adt-3* | *adt-3 promoter 1::rfp* | 181 | 1 | +4517 |
| *R03E9.2* | *R03E9.2 promoter 1::rfp* | 172 | 2 | −5259, −5196 |
|  | *R03E9.2 promoter 2::rfp* | 182 | 1 | −5050 |
|  | *R03E9.2 promoter 3::rfp* | 158 | 1 | −3428 |
|  | *R03E9.2 promoter 4::rfp* | 170 | 1 | −2180 |
| *F49B2.6* | *F49B2.6 promoter 1::rfp* | 153 | 1 | −1459 |
|  | *F49B2.6 promoter 2::rfp* | 154 | 2 | −198, −154 |
| *nta-1* | *nta-1 promoter 1::rfp* | 156 | 1 | −69 |
|  | *nta-1 promoter 2::rfp* | 154 | 1 | +145 |
| *T14B4.9* | *T14B4.9 promoter 1::gfp* | 172 | 1 | −1647 |
|  | *T14B4.9 promoter 2::gfp* | 254 | 2 | −393, −279 |
|  | *T14B4.9 promoter 3::gfp* | 171 | 1 | +210 |
| *tbc-12* | *tbc-12 promoter 1::gfp* | 172 | 2 | +1481, +1545 |
|  | *tbc-12 promoter 2::gfp* | 155 | 1 | +1669 |
|  | *tbc-12 promoter 3::gfp* | 232 | 2 | +2086, +2208 |
|  | *tbc-12 promoter 4::gfp* | 232 | 1 | +2918 |
| *pgp-4* | *pgp-4 promoter 1::rfp* | 187 | 1 | −325 |
|  | *pgp-4 promoter 2::rfp* | 157 | 1 | +558 |
| *haf-7* | *haf-7 promoter 1::rfp* | 178 | 2 | −1468, −1370 |
|  | *haf-7 promoter 2::rfp* | 204 | 1 | +2134 |

1. Slattery, M., Zhou, T., Yang, L., Dantas Machado, A.C., Gordan, R., and Rohs, R. (2014). Absence of a simple code: how transcription factors read the genome. Trends Biochem Sci 39, 381-399. 10.1016/j.tibs.2014.07.002.

## APPENDIX

**Appendix Figure S1.**
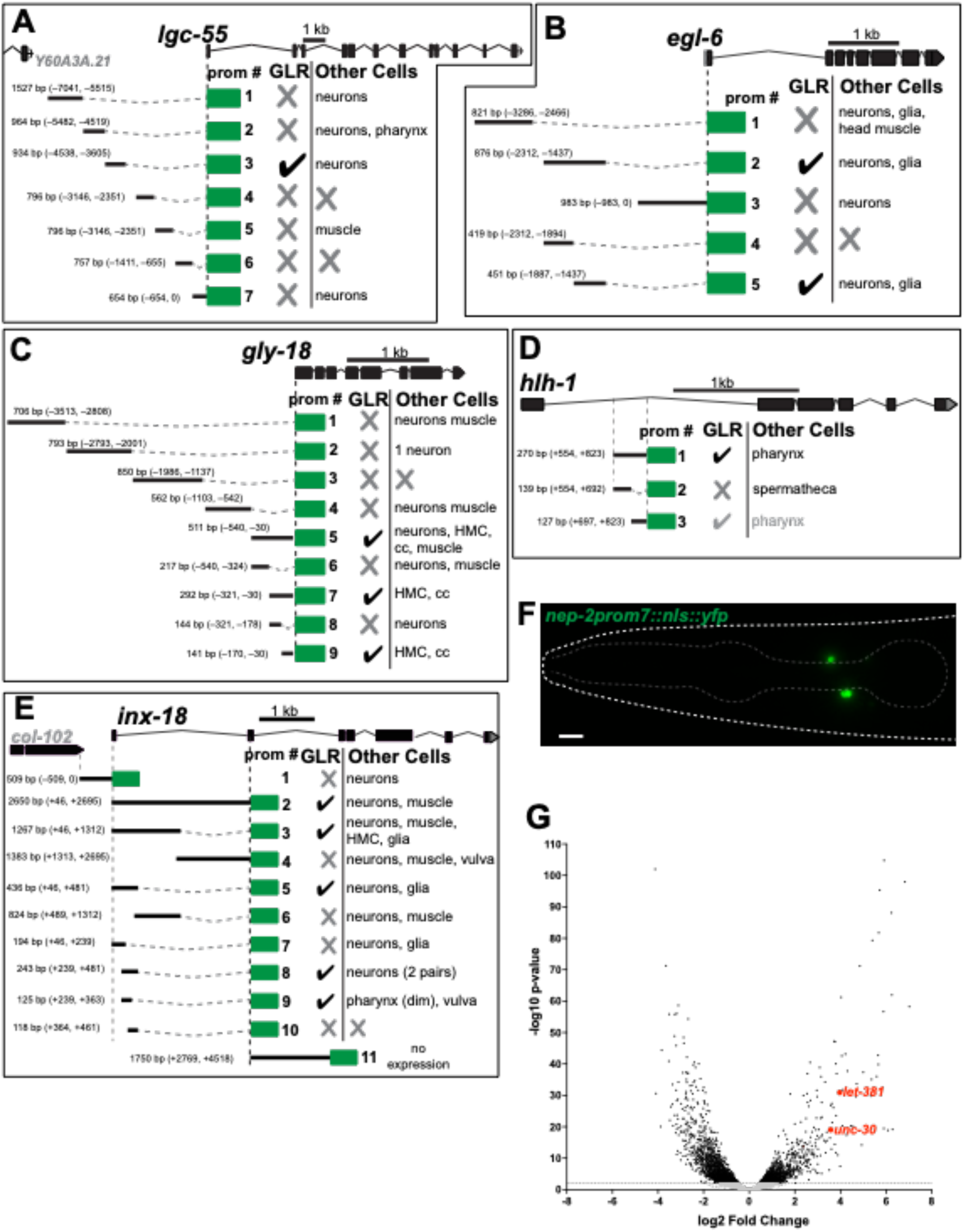
Analysis of cis-regulatory regions of GLR glia-expressed genes. (A) – (E) Cis-regulatory dissection analysis for the genes *lgc-55* (A), *egl-6* (B), *gly-18* (C), *hlh-1* (F) and *inx-18* (E). (F) A stable transgenic strain expressing nuclear YFP under the GLR glia-specific *nep-2prom7* was used to isolate GLR glia for the transcriptome analysis. Anterior is left, dorsal is up and scale bar is 10 μm. (G) Volcano plot show significantly enriched and depleted genes (p-value >0.05) among all GLR glia expressed genes (>50 reads). *let-381* and *unc-30*, the two transcription factor genes we show to have important roles in GLR glia development, are among the most highly enriched genes.

**Appendix Figure S2.**
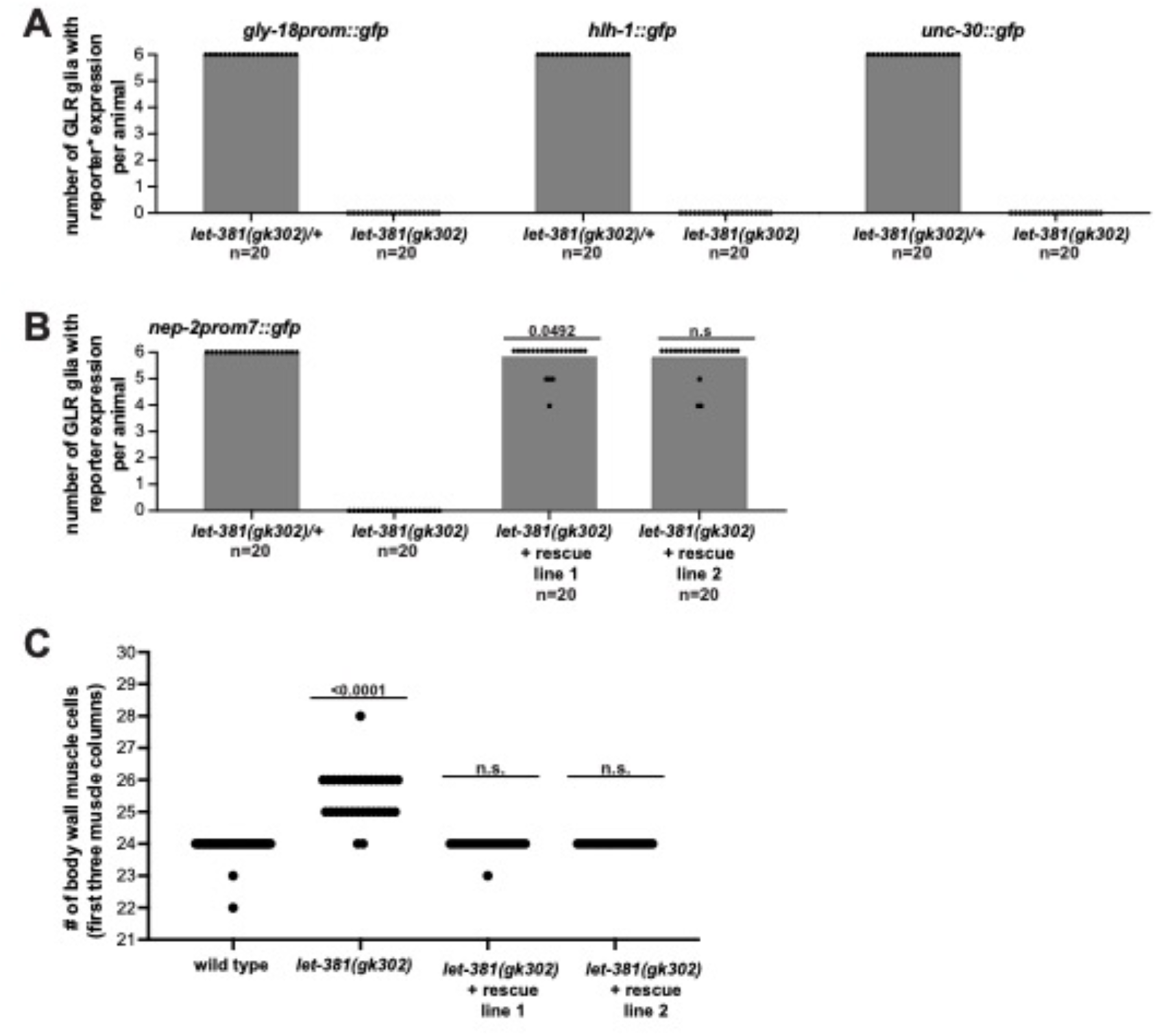
*let-381/FoxF* is required for GLR glia fate specification. (A) Number of GLR glia expressing *gly-18prom::gfp*, *hlh-1::gfp* and *unc-30::gfp* expression in wild type and homozygous *let-381(gk302)* mutants. Expression of all genes tested is absent from GLR cells in *let-381* homozygous null mutants. (B), (C) Animals transgenic (2 independent lines) for the fosmid genomic clone WRM069bF08 carrying wild type *let-381* have (B) 6 GLR glia and (C) no extra head muscle cells. Data information: Un-paired t-test used for statistical analysis in (B) and (C).

**Appendix Figure Fig. S3.**
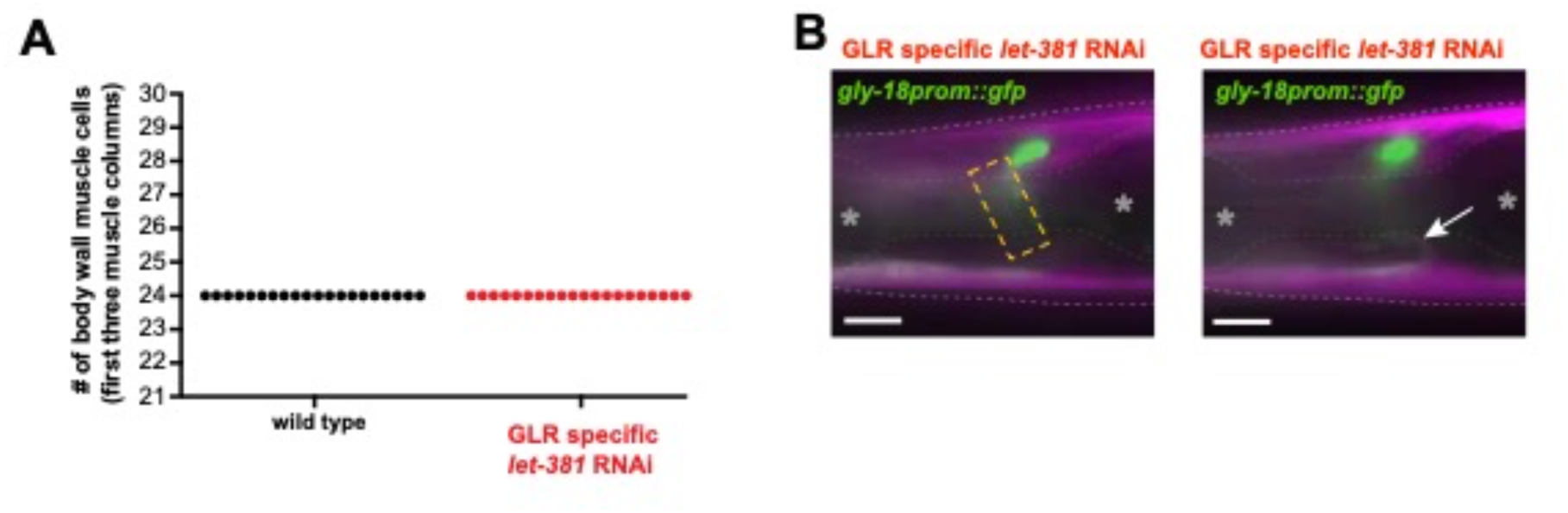
Postembryonic *let-381* knockdown does not affect GLR fate specification. (A) Animals carrying the GLR glia-specific *let-381* RNAi arrays do not have extra head muscle cells. (B) In addition, *let-381* RNAi does not cause anteriorly displaced Nerve Ring. Yellow dashed box outlines the Nerve Ring (green) on the left image, and arrow on the right image shows a head muscle arm extending into the Nerve Ring. As evidenced NR is located in its normal position between the two pharyngeal bulbs (grey asterisks). Data information: Anterior is left, dorsal is up and scale bars are 10 μm for all animal images.

## REFERENCES

1. Allen, N.J., and Lyons, D.A. (2018). Glia as architects of central nervous system formation and function. Science 362, 181–185. 10.1126/science.aat0473.

2. Rowitch, D.H., and Kriegstein, A.R. (2010). Developmental genetics of vertebrate glial-cell specification. Nature 468, 214–222. 10.1038/nature09611.

3. Kastriti, M.E., and Adameyko, I. (2017). Specification, plasticity and evolutionary origin of peripheral glial cells. Current Opinion in Neurobiology 47, 196–202. 10.1016/j.conb.2017.11.004.

4. Ginhoux, F., and Prinz, M. (2015). Origin of microglia: current concepts and past controversies. Cold Spring Harb Perspect Biol 7, a020537. 10.1101/cshperspect.a020537.

5. Ginhoux, F., Greter, M., Leboeuf, M., Nandi, S., See, P., Gokhan, S., Mehler, M.F., Conway, S.J., Ng, L.G., Stanley, E.R., et al. (2010). Fate mapping analysis reveals that adult microglia derive from primitive macrophages. Science 330, 841–845. 10.1126/science.1194637.

6. Ginhoux, F., Lim, S., Hoeffel, G., Low, D., and Huber, T. (2013). Origin and differentiation of microglia. Front Cell Neurosci 7, 45. 10.3389/fncel.2013.00045.

7. Wegner, M. (2020). Chapter 34 - Specification of oligodendrocytes. In Patterning and Cell Type Specification in the Developing CNS and PNS (Second Edition), J. Rubenstein, P. Rakic, B. Chen, and K.Y. Kwan, eds. (Academic Press), pp. 847–866. 10.1016/B978-0-12-814405-3.00034-5.

8. Hochstim, C., Deneen, B., Lukaszewicz, A., Zhou, Q., and Anderson, D.J. (2008). Identification of Positionally Distinct Astrocyte Subtypes whose Identities Are Specified by a Homeodomain Code. Cell 133, 510–522. 10.1016/j.cell.2008.02.046.

9. Canals, I., Ginisty, A., Quist, E., Timmerman, R., Fritze, J., Miskinyte, G., Monni, E., Hansen, M.G., Hidalgo, I., Bryder, D., et al. (2018). Rapid and efficient induction of functional astrocytes from human pluripotent stem cells. Nature Methods 15, 693–696. 10.1038/s41592-018-0103-2.

10. Shaham, S. (2015). Glial development and function in the nervous system of Caenorhabditis elegans. Cold Spring Harb Perspect Biol 7, a020578. 10.1101/cshperspect.a020578.

11. Sulston, J.E., Schierenberg, E., White, J.G., and Thomson, J.N. (1983). The embryonic cell lineage of the nematode Caenorhabditis elegans. Developmental Biology 100, 64–119. 10.1016/0012-1606(83)90201-4.

12. Shaham, S. (2015). Glial development and function in the nervous system of Caenorhabditis elegans. Cold Spring Harb. Perspect 8, 4.

13. Wallace, Sean W., Singhvi, A., Liang, Y., Lu, Y., and Shaham, S. (2016). PROS-1/Prospero Is a Major Regulator of the Glia-Specific Secretome Controlling Sensory-Neuron Shape and Function in C. elegans. Cell Reports 15, 550–562. 10.1016/j.celrep.2016.03.051.

14. Zhang, A., Noma, K., and Yan, D. (2020). Regulation of Gliogenesis by lin-32/Atoh1 in Caenorhabditis elegans. G3 Genes|Genomes|Genetics 10, 3271–3278. 10.1534/g3.120.401547.

15. Mizeracka, K., Rogers, J.M., Rumley, J.D., Shaham, S., Bulyk, M.L., Murray, J.I., and Heiman, M.G. (2021). Lineage-specific control of convergent differentiation by a Forkhead repressor. Development 148. 10.1242/dev.199493.

16. White, J.G., Southgate, E., Thomson, J.N., and Brenner, S. (1986). The structure of the nervous system of the nematode Caenorhabditis elegans. Philos Trans R Soc Lond B Biol Sci 314, 1–340.

17. Gendrel, M., Atlas, E.G., and Hobert, O. (2016). A cellular and regulatory map of the GABAergic nervous system of C. elegans. Elife 5. 10.7554/eLife.17686.

18. Nass, R., Hall, D.H., Miller, D.M., 3rd, and Blakely, R.D. (2002). Neurotoxin-induced degeneration of dopamine neurons in Caenorhabditis elegans. Proc Natl Acad Sci U S A 99, 3264–3269. 10.1073/pnas.042497999.

19. Altun, Z.F., and Hall, D.H. (2016). Handbook of *C. elegans* Anatomy. In WormAtlas.

20. Yamada, K., Hirotsu, T., Matsuki, M., Butcher, R.A., Tomioka, M., Ishihara, T., Clardy, J., Kunitomo, H., and Iino, Y. (2010). Olfactory plasticity is regulated by pheromonal signaling in Caenorhabditis elegans. Science 329, 1647–1650.

21. Ringstad, N., Abe, N., and Horvitz, H.R. (2009). Ligand-gated chloride channels are receptors for biogenic amines in C. elegans. Science 325, 96–100.

22. Ringstad, N., and Horvitz, H.R. (2008). FMRFamide neuropeptides and acetylcholine synergistically inhibit egg-laying by C. elegans. Nature neuroscience 11, 1168–1176.

23. Warren, C.E., Krizus, A., and Dennis, J.W. (2001). Complementary expression patterns of six nonessential Caenorhabditis elegans core 2/I N-acetylglucosaminyltransferase homologues. Glycobiology 11, 979–988.

24. Krause, M., Harrison, S.W., Xu, S.-Q., Chen, L., and Fire, A. (1994). Elements regulating cell-and stage-specific expression of the C. elegans MyoD family homolog hlh-1. Developmental biology 166, 133–148.

25. Mi, H., Muruganujan, A., Huang, X., Ebert, D., Mills, C., Guo, X., and Thomas, P.D. (2019). Protocol Update for large-scale genome and gene function analysis with the PANTHER classification system (v.14.0). Nat Protoc 14, 703–721. 10.1038/s41596-019-0128-8.

26. Thomas, P.D., Ebert, D., Muruganujan, A., Mushayahama, T., Albou, L.P., and Mi, H. (2022). PANTHER: Making genome-scale phylogenetics accessible to all. Protein Sci 31, 8–22. 10.1002/pro.4218.

27. Yang, Y., and Jackson, R. (2019). Astrocyte identity: evolutionary perspectives on astrocyte functions and heterogeneity. Current Opinion in Neurobiology 56, 40–46. 10.1016/j.conb.2018.11.006.

28. Zhang, Y., Chen, K., Sloan, S.A., Bennett, M.L., Scholze, A.R., O’Keeffe, S., Phatnani, H.P., Guarnieri, P., Caneda, C., Ruderisch, N., et al. (2014). An RNA-Sequencing Transcriptome and Splicing Database of Glia, Neurons, and Vascular Cells of the Cerebral Cortex. The Journal of Neuroscience 34, 11929–11947. 10.1523/jneurosci.1860-14.2014.

29. Batiuk, M.Y., Martirosyan, A., Wahis, J., de Vin, F., Marneffe, C., Kusserow, C., Koeppen, J., Viana, J.F., Oliveira, J.F., Voet, T., et al. (2020). Identification of region-specific astrocyte subtypes at single cell resolution. Nature Communications 11, 1220. 10.1038/s41467-019-14198-8.

30. Munji, R.N., Soung, A.L., Weiner, G.A., Sohet, F., Semple, B.D., Trivedi, A., Gimlin, K., Kotoda, M., Korai, M., Aydin, S., et al. (2019). Profiling the mouse brain endothelial transcriptome in health and disease models reveals a core blood–brain barrier dysfunction module. Nature Neuroscience 22, 1892–1902. 10.1038/s41593-019-0497-x.

31. Amin, N.M., Shi, H., and Liu, J. (2010). The FoxF/FoxC factor LET-381 directly regulates both cell fate specification and cell differentiation in C. elegans mesoderm development. Development 137, 1451–1460. 10.1242/dev.048496.

32. Scimone, M.L., Wurtzel, O., Malecek, K., Fincher, C.T., Oderberg, I.M., Kravarik, K.M., and Reddien, P.W. (2018). foxF-1 Controls Specification of Non-body Wall Muscle and Phagocytic Cells in Planarians. Curr Biol 28, 3787–3801 e3786. 10.1016/j.cub.2018.10.030.

33. Reyahi, A., Nik, A.M., Ghiami, M., Gritli-Linde, A., Ponten, F., Johansson, B.R., and Carlsson, P. (2015). Foxf2 Is Required for Brain Pericyte Differentiation and Development and Maintenance of the Blood-Brain Barrier. Dev Cell 34, 19–32. 10.1016/j.devcel.2015.05.008.

34. Shah, P.K., Santella, A., Jacobo, A., Siletti, K., Hudspeth, A.J., and Bao, Z. (2017). An In Toto Approach to Dissecting Cellular Interactions in Complex Tissues. Dev Cell 43, 530–540 e534. 10.1016/j.devcel.2017.10.021.

35. Esposito, G., Di Schiavi, E., Bergamasco, C., and Bazzicalupo, P. (2007). Efficient and cell specific knock-down of gene function in targeted C. elegans neurons. Gene 395, 170–176. 10.1016/j.gene.2007.03.002.

36. Zhang, L., Ward, J.D., Cheng, Z., and Dernburg, A.F. (2015). The auxin-inducible degradation (AID) system enables versatile conditional protein depletion in C. elegans. Development 142, 4374–4384. 10.1242/dev.129635.

37. Martinez, M.A.Q., Kinney, B.A., Medwig-Kinney, T.N., Ashley, G., Ragle, J.M., Johnson, L., Aguilera, J., Hammell, C.M., Ward, J.D., and Matus, D.Q. (2020). Rapid Degradation of Caenorhabditis elegans Proteins at Single-Cell Resolution with a Synthetic Auxin. G3 (Bethesda) 10, 267–280. 10.1534/g3.119.400781.

38. Hobert, O. (2011). Regulation of terminal differentiation programs in the nervous system. Annu Rev Cell Dev Biol 27, 681–696. 10.1146/annurev-cellbio-092910-154226.

39. Bailey, T.L., and Elkan, C. (1994). Fitting a mixture model by expectation maximization to discover motifs in biopolymers. Proc Int Conf Intell Syst Mol Biol 2, 28–36.

40. Peterson, R.S., Lim, L., Ye, H., Zhou, H., Overdier, D.G., and Costa, R.H. (1997). The winged helix transcriptional activator HFH-8 is expressed in the mesoderm of the primitive streak stage of mouse embryos and its cellular derivatives. Mech Dev 69, 53–69. 10.1016/s0925-4773(97)00153-6.

41. Narasimhan, K., Lambert, S.A., Yang, A.W., Riddell, J., Mnaimneh, S., Zheng, H., Albu, M., Najafabadi, H.S., Reece-Hoyes, J.S., Fuxman Bass, J.I., et al. (2015). Mapping and analysis of Caenorhabditis elegans transcription factor sequence specificities. Elife 4. 10.7554/eLife.06967.

42. Jin, Y., Hoskins, R., and Horvitz, H.R. (1994). Control of type-D GABAergic neuron differentiation by C. elegans UNC-30 homeodomain protein. Nature 372, 780–783. 10.1038/372780a0.

43. Eastman, C., Horvitz, H.R., and Jin, Y. (1999). Coordinated transcriptional regulation of the unc-25 glutamic acid decarboxylase and the unc-47 GABA vesicular transporter by the Caenorhabditis elegans UNC-30 homeodomain protein. J Neurosci 19, 6225–6234. 10.1523/JNEUROSCI.19-15-06225.1999.

44. Cinar, H., Keles, S., and Jin, Y. (2005). Expression profiling of GABAergic motor neurons in Caenorhabditis elegans. Curr Biol 15, 340–346. 10.1016/j.cub.2005.02.025.

45. Bayer, E.A., Stecky, R.C., Neal, L., Katsamba, P.S., Ahlsen, G., Balaji, V., Hoppe, T., Shapiro, L., Oren-Suissa, M., and Hobert, O. (2020). Ubiquitin-dependent regulation of a conserved DMRT protein controls sexually dimorphic synaptic connectivity and behavior. Elife 9. 10.7554/eLife.59614.

46. Katz, M., Corson, F., Iwanir, S., Biron, D., and Shaham, S. (2018). Glia Modulate a Neuronal Circuit for Locomotion Suppression during Sleep in C. elegans. Cell Rep 22, 2575–2583. 10.1016/j.celrep.2018.02.036.

47. Katz, M., Corson, F., Keil, W., Singhal, A., Bae, A., Lu, Y., Liang, Y., and Shaham, S. (2019). Glutamate spillover in C. elegans triggers repetitive behavior through presynaptic activation of MGL-2/mGluR5. Nat Commun 10, 1882. 10.1038/s41467-019-09581-4.

48. Zallen, J.A., Kirch, S.A., and Bargmann, C.I. (1999). Genes required for axon pathfinding and extension in the C. elegans nerve ring. Development 126, 3679–3692. 10.1242/dev.126.16.3679.

49. Kennerdell, J.R., Fetter, R.D., and Bargmann, C.I. (2009). Wnt-Ror signaling to SIA and SIB neurons directs anterior axon guidance and nerve ring placement in C. elegans. Development 136, 3801–3810. 10.1242/dev.038109.

50. Choi, U., Hu, M., and Sieburth, D. (2022). hmc, a cell with previously unknown function couples neuropeptide transmitters with muscle contraction during a rhythmic behavior in C. elegans. Research Square.

51. Hobert, O., and Kratsios, P. (2019). Neuronal identity control by terminal selectors in worms, flies, and chordates. Curr Opin Neurobiol 56, 97–105. 10.1016/j.conb.2018.12.006.

52. Remesal, L., Roger-Baynat, I., Chirivella, L., Maicas, M., Brocal-Ruiz, R., Perez-Villalba, A., Cucarella, C., Casado, M., and Flames, N. (2020). PBX1 acts as terminal selector for olfactory bulb dopaminergic neurons. Development 147. 10.1242/dev.186841.

53. Feng, W., Li, Y., Dao, P., Aburas, J., Islam, P., Elbaz, B., Kolarzyk, A., Brown, A.E., and Kratsios, P. (2020). A terminal selector prevents a Hox transcriptional switch to safeguard motor neuron identity throughout life. Elife 9. 10.7554/eLife.50065.

54. Zaffran, S., Kuchler, A., Lee, H.H., and Frasch, M. (2001). biniou (FoxF), a central component in a regulatory network controlling visceral mesoderm development and midgut morphogenesis in Drosophila. Genes Dev 15, 2900–2915. 10.1101/gad.917101.

55. Mahlapuu, M., Ormestad, M., Enerback, S., and Carlsson, P. (2001). The forkhead transcription factor Foxf1 is required for differentiation of extra-embryonic and lateral plate mesoderm. Development 128, 155–166. 10.1242/dev.128.2.155.

56. Tseng, H.T., Shah, R., and Jamrich, M. (2004). Function and regulation of FoxF1 during Xenopus gut development. Development 131, 3637–3647. 10.1242/dev.01234.

57. Ormestad, M., Astorga, J., Landgren, H., Wang, T., Johansson, B.R., Miura, N., and Carlsson, P. (2006). Foxf1 and Foxf2 control murine gut development by limiting mesenchymal Wnt signaling and promoting extracellular matrix production. Development 133, 833–843. 10.1242/dev.02252.

58. Saunders, A., Macosko, E.Z., Wysoker, A., Goldman, M., Krienen, F.M., de Rivera, H., Bien, E., Baum, M., Bortolin, L., Wang, S., et al. (2018). Molecular Diversity and Specializations among the Cells of the Adult Mouse Brain. Cell 174, 1015–1030.e1016. 10.1016/j.cell.2018.07.028.

59. Vanlandewijck, M., He, L., Mäe, M.A., Andrae, J., Ando, K., Del Gaudio, F., Nahar, K., Lebouvier, T., Laviña, B., Gouveia, L., et al. (2018). A molecular atlas of cell types and zonation in the brain vasculature. Nature 554, 475–480. 10.1038/nature25739.

60. Hupe, M., Li, M.X., Kneitz, S., Davydova, D., Yokota, C., Kele, J., Hot, B., Stenman, J.M., and Gessler, M. (2017). Gene expression profiles of brain endothelial cells during embryonic development at bulk and single-cell levels. Sci Signal 10. 10.1126/scisignal.aag2476.

61. Kalinichenko, V.V., Gusarova, G.A., Shin, B., and Costa, R.H. (2003). The forkhead box F1 transcription factor is expressed in brain and head mesenchyme during mouse embryonic development. Gene Expr Patterns 3, 153–158. 10.1016/s1567-133x(03)00010-3.

62. Bundgaard, M., and Abbott, N.J. (2008). All vertebrates started out with a glial blood-brain barrier 4-500 million years ago. Glia 56, 699–708. 10.1002/glia.20642.

63. Brenner, S. (1974). The genetics of Caenorhabditis elegans. Genetics 77, 71–94.

64. Dokshin, G.A., Ghanta, K.S., Piscopo, K.M., and Mello, C.C. (2018). Robust Genome Editing with Short Single-Stranded and Long, Partially Single-Stranded DNA Donors in Caenorhabditis elegans. Genetics 210, 781–787. 10.1534/genetics.118.301532.

65. Goudeau, J., Sharp, C.S., Paw, J., Savy, L., Leonetti, M.D., York, A.G., Updike, D.L., Kenyon, C., and Ingaramo, M. (2021). Split-wrmScarlet and split-sfGFP: tools for faster, easier fluorescent labeling of endogenous proteins in Caenorhabditis elegans. Genetics 217. 10.1093/genetics/iyab014.

66. Hobert, O. (2002). PCR fusion-based approach to create reporter gene constructs for expression analysis in transgenic C. elegans. Biotechniques 32, 728–730. 10.2144/02324bm01.

67. Bacaj, T., Tevlin, M., Lu, Y., and Shaham, S. (2008). Glia are essential for sensory organ function in C. elegans. Science 322, 744–747.

68. Kage-Nakadai, E., Imae, R., Yoshina, S., and Mitani, S. (2014). Methods for single/low-copy integration by ultraviolet and trimethylpsoralen treatment in Caenorhabditis elegans. Methods 68, 397–402. 10.1016/j.ymeth.2014.02.036.

69. Tursun, B., Cochella, L., Carrera, I., and Hobert, O. (2009). A toolkit and robust pipeline for the generation of fosmid-based reporter genes in C. elegans. PLoS One 4, e4625. 10.1371/journal.pone.0004625.

70. Stefanakis, N., Carrera, I., and Hobert, O. (2015). Regulatory Logic of Pan-Neuronal Gene Expression in C. elegans. Neuron 87, 733–750. 10.1016/j.neuron.2015.07.031.

71. Kennedy, S., Wang, D., and Ruvkun, G. (2004). A conserved siRNA-degrading RNase negatively regulates RNA interference in C. elegans. Nature 427, 645–649. 10.1038/nature02302.

72. Dobin, A., Davis, C.A., Schlesinger, F., Drenkow, J., Zaleski, C., Jha, S., Batut, P., Chaisson, M., and Gingeras, T.R. (2012). STAR: ultrafast universal RNA-seq aligner. Bioinformatics 29, 15–21. 10.1093/bioinformatics/bts635.

73. DeLuca, D.S., Levin, J.Z., Sivachenko, A., Fennell, T., Nazaire, M.D., Williams, C., Reich, M., Winckler, W., and Getz, G. (2012). RNA-SeQC: RNA-seq metrics for quality control and process optimization. Bioinformatics 28, 1530–1532. 10.1093/bioinformatics/bts196.

74. Liao, Y., Smyth, G.K., and Shi, W. (2013). featureCounts: an efficient general purpose program for assigning sequence reads to genomic features. Bioinformatics 30, 923–930. 10.1093/bioinformatics/btt656.

75. Love, M.I., Huber, W., and Anders, S. (2014). Moderated estimation of fold change and dispersion for RNA-seq data with DESeq2. Genome Biology 15, 550. 10.1186/s13059-014-0550-8.

76. Edelstein, A., Amodaj, N., Hoover, K., Vale, R., and Stuurman, N. (2010). Computer control of microscopes using µManager. Curr Protoc Mol Biol Chapter 14, Unit14.20. 10.1002/0471142727.mb1420s92.

77. Schneider, C.A., Rasband, W.S., and Eliceiri, K.W. (2012). NIH Image to ImageJ: 25 years of image analysis. Nature Methods 9, 671–675. 10.1038/nmeth.2089.

78. large-scale screening for targeted knockouts in the Caenorhabditis elegans genome. (2012). G3 (Bethesda) 2, 1415–1425. 10.1534/g3.112.003830.

79. Howell, A.M., Gilmour, S.G., Mancebo, R.A., and Rose, A.M. (1987). Genetic analysis of a large autosomal region in Caenorhabditis elegans by the use of a free duplication. Genetics Research 49, 207–213. 10.1017/S0016672300027099.

80. Forrester, W.C., Dell, M., Perens, E., and Garriga, G. (1999). A C. elegans Ror receptor tyrosine kinase regulates cell motility and asymmetric cell division. Nature 400, 881–885. 10.1038/23722.

81. Kostas, S.A., and Fire, A. (2002). The T-box factor MLS-1 acts as a molecular switch during specification of nonstriated muscle in C. elegans. Genes Dev 16, 257–269. 10.1101/gad.923102.

82. Liu, X., Long, F., Peng, H., Aerni, S.J., Jiang, M., Sánchez-Blanco, A., Murray, J.I., Preston, E., Mericle, B., Batzoglou, S., et al. (2009). Analysis of Cell Fate from Single-Cell Gene Expression Profiles in C. elegans. Cell 139, 623–633. 10.1016/j.cell.2009.08.044.

83. Sarov, M., Schneider, S., Pozniakovski, A., Roguev, A., Ernst, S., Zhang, Y., Hyman, A.A., and Stewart, A.F. (2006). A recombineering pipeline for functional genomics applied to Caenorhabditis elegans. Nat Methods 3, 839–844. 10.1038/nmeth933.

84. McKay, S.J., Johnsen, R., Khattra, J., Asano, J., Baillie, D.L., Chan, S., Dube, N., Fang, L., Goszczynski, B., Ha, E., et al. (2003). Gene expression profiling of cells, tissues, and developmental stages of the nematode C. elegans. Cold Spring Harb Symp Quant Biol 68, 159–169. 10.1101/sqb.2003.68.159.

85. Altun-Gultekin, Z., Andachi, Y., Tsalik, E.L., Pilgrim, D., Kohara, Y., and Hobert, O. (2001). A regulatory cascade of three homeobox genes, ceh-10, ttx-3 and ceh-23, controls cell fate specification of a defined interneuron class in C. elegans. Development 128, 1951–1969. 10.1242/dev.128.11.1951.

86. Slattery, M., Zhou, T., Yang, L., Dantas Machado, A.C., Gordan, R., and Rohs, R. (2014). Absence of a simple code: how transcription factors read the genome. Trends Biochem Sci 39, 381–399. 10.1016/j.tibs.2014.07.002.

